# On the shuttling across the blood-brain barrier via tubules formation: mechanism and cargo avidity bias

**DOI:** 10.1101/2020.04.04.025866

**Authors:** Xiaohe Tian, Diana Moreira Leite, Edoardo Scarpa, Sophie Nyberg, Gavin Fullstone, Joe Forth, Diana Lourenço Matias, Azzurra Apriceno, Alessandro Poma, Aroa Duro-Castano, Manish Vuyyuru, Lena Harker-Kirschneck, Anđela Šarić, Zhongping Zhang, Pan Xiang, Bin Fang, Yupeng Tian, Lei Luo, Loris Rizzello, Giuseppe Battaglia

## Abstract

The blood-brain barrier is made of polarised brain endothelial cells (BECs) phenotypically conditioned by the central nervous system (CNS). Transport across BECs is of paramount importance for nutrient uptake as well as to rid the brain of waste products. Nevertheless, currently we do not understand how large macromolecular cargo shuttles across and how BECs discriminate between the brain-bound and own nutrients. Here, we study the low-density lipoprotein receptor-related protein 1 (LRP1) an essential regulator of BEC transport, and show that it is associated with endocytic effectors, endo-lysosomal compartments as well as syndapin-2, a member of the Bin/Amphiphysin/Rvs (BAR) domain superfamily known to stabilise tubular carriers. We employed synthetic self-assembled vesicles, polymersomes, as a multivalent system with tunable avidity as a tool to investigate the mechanism of transport across BECs. We used a combination of conventional and super-resolution microscopy, both *in vivo* and *in vitro*, accompanied with biophysical modelling of transport kinetics and membrane-bound interactions. Our results demonstrate that the avidity of the ligand-receptor interaction (the overall cargo binding energy) determines the mechanism of sorting during the early stages of endocytosis and consequent trafficking. We show that high avidity cargo biases the LRP1 towards internalisation and fast degradation in BECs, while mid avidity augments the formation of syndapin-2 stabilised tubular carriers and promotes fast shuttling across BECs. Thus, we map out a very detailed mechanism where clathrin, actin, syndapin-2, dynamin and SNARE act synergistically to enable fast shuttling across BECs.

## Introduction

The human brain accounts for about 2-3% of the total body mass, and yet it consumes up to 50% of the total intake of oxygen and glucose [1]. Such a high energy demand is only possible because of a controlled gating of mass exchange with the body across a network of barriers that are phenotypically regulated by the brain cells. The most important of all gateways is the blood-brain barrier (BBB). This is richest capillary network in the body being able to feed the brain components very effectively with about one capillary *per* neuron and about 10 to 15 *μ*m of the average distance between one capillary to another [2, 3, 4]. Capillaries are made by polarised endothelial cells connected via tight junctions. Brain endothelial cells (BECs) are conditioned by the neighbouring brain cells to limit passive transport by forming impermeable tight junctions, lacking fenestrations and expressing efflux transporters that protect the brain from harmful compounds [5, 3, 4]. BBB dysfunctions are at the core of ageing, neurological degeneration, stroke, and multiple sclerosis [5]. Notably, the BBB makes the brain impermeable to most therapeutics, leading to a bottleneck in drug development [4].

BECs control the transport of small molecules, such as glucose and amino acids, by expressing specialised solute carrier transporters on the apical (blood) and basal (brain) membranes that pump molecules across one-by-one [6]. BECs overexpress transferrin [7], insulin receptors [8], and low-density lipoprotein receptor-related protein 1 (LRP1) [9, 10, 11] and these receptors are often involved in shuttling their respective ligands into trafficking membrane-enveloped carriers across the cell *via* a process collectively known as transcytosis [12, 6]. LRP1 is a critical motif highly expressed by neurons [13] and astrocytes [14], and it has been reported to bind more than 40 ligands [10] undergoing rapid endocytosis with a half-life of less than 30 seconds [15, 10]. The functions of LRP1 include the control of trafficking and regulation of several intracellular signalling pathways [16] making its deletion in mice lethal [17]. At the BBB, tissue-type plasminogen activator binding to LRP1 triggers its cleavage with consequences on the BBB integrity [18]. LRP1 has been singled out for the BBB brain-to-blood efflux of secreted amyloid precursor protein (APP) containing Kunitz protease inhibitor (KPI) domain [19]. In Alzheimer’s disease, LRP1 is an important regulator [20] with patients showing lower receptor levels [21] and with localised LRP1 deletion leading to amyloid-*β* retention and cognition impairment. Most relevant to the present work, LRP1 has also been associated with the blood-to-brain efflux of lactoferrin [22], receptor-associated protein (RAP) [23] and KPI domain-containing proteins [24].

Transcytosis is an active transport involving the rearrangement of large membrane volumes, and although it has been studied in detail in other barrier tissues (such as the epithelium) little is known about it in endothelial cells [12, 8, 25]. Epithelial, and by analogy endothelial, transcytosis involves three steps: (i) endocytosis; a vesicular carrier emerges from one side of the membrane, typically involving clathrin or caveolin, (ii) trafficking; where the carrier moves towards and fuses with the endo-lysosome network and eventually (iii) exocytosis; where a new vesicular carrier emerges from endo-lysosome, move towards and fuse with opposite side of the plasma membrane [8]. Such a sequence of events is viable in thick epithelial cells but often endothelium can be as thin as few hundreds of nanometres [25], and as such the internal volume is too small to house the machinery associated with the three transcytosis steps. In addition to the traditional spherical trafficking vesicles, ultra-structural observations revealed the formation of “pores” or “channels” spanning the endothelial cell referred to as transendothelial channels (TEC) [26]. Bundgaard reconstructed 3D projections from serial sections of transmission electron micrographs of hagfish BECs showing that intracellular membranes arising from transcytosis were rarely single vesicles, but instead part of large multidimensional dendritic networks or ‘tubes’ [27]. Tubular networks and chains of vesiculo-vacuolar organelles (VVOs) were also reported in fenestrated endothelium [28]. Despite being widely observed using electron microscopy, the molecular identity and the mechanism regulating the formation of both TECs and VVOs is still poorly defined. Although early hypotheses supported the idea that VVOs were clusters of caveolin-rich vesicles, further investigations demonstrated the presence of VVOs even after the knock-down of caveolin [29]. Recently, caveolin lack of involvement was further demonstrated in BECs showing that a unique BBB marker, the MFSD2a [30, 31], inhibits the formation of caveolae [32]. A piece of critical information missing from all these studies is the role of membrane sculpting proteins, most notably those comprising the Bin/Amphiphysin/Rvs (BAR) domain [33]. These are critical scaffolds that bind, polymerise onto curved membranes, and link with essential components such as actin and dynamin [33]. Among the different types of BAR proteins, the Fer-CIP4 homology-BAR (F-BAR) family [34], and syndapin-2 in particular possess a unique ability in creating membrane tubules [35].

Herein, we elucidate the trafficking of LRP1 in BECs and correlate its transcytosis mechanism with syndapin-2 using both *in vitro* and *in vivo* models of the BBB. We use synthetic vesicles, polymersomes (POs), functionalised with LRP1 targeting moieties established to traverse the BBB both in mice [36] and rats [37], to assess how multivalency and hence binding avidity control LRP1-mediated transcytosis.

## Results and Discussion

### LRP1 trafficking across brain endothelium

In order to study the subcellular localisation of LRP1 associated proteins in BECs, we employed a well-established 3D model of the BBB consisting of confluent mouse brain endothelioma cells (bEnd3) cultured onto collagen-coated porous transwell inserts (**Fig.S1a**). We have established the barrier properties of this BBB model [36, 38] by measuring both the transendothelial resistance and an apparent permeability coefficient of different molecular mass dextran (*P*), calculated as:

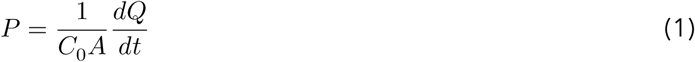

with *C*_0_ being the initial cargo concentration, *A* the total surface area of the transwell membrane, and 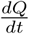 the transport rate calculated as the gradient of mass over time. For 4 and 70 kDa dextrans we measured a permeability of *P*_4*kDa*_ = 19.6 and *P*_70*kDa*_ = 4.7 nm s^−1^, respectively (**Fig.S1b**). bEnd3 cell monolayer presented a classical morphology with the expression of platelet endothelial cell adhesion molecule (PECAM-1), and tight junction proteins, claudin-5 and zona occludens 1 (ZO-1) (**Fig.S1c**). Most relevant to the present study, we confirmed the expression of LRP1 in BECs using both western blot (WB) (**Fig.1a**) and immuno-fluorescence (**Fig.1b**) targeting the cytosolic and extracellular domains, respectively. The micrographs collected across different monolayer regions show the wide expression of LRP1 in BECs (**Fig.1b, c**). Moreover, 3D reconstructions evidence that LRP1 is expressed on both the apical and basal cell surfaces as well as in the perinuclear area (**Fig.1d**).

**Figure 1:**
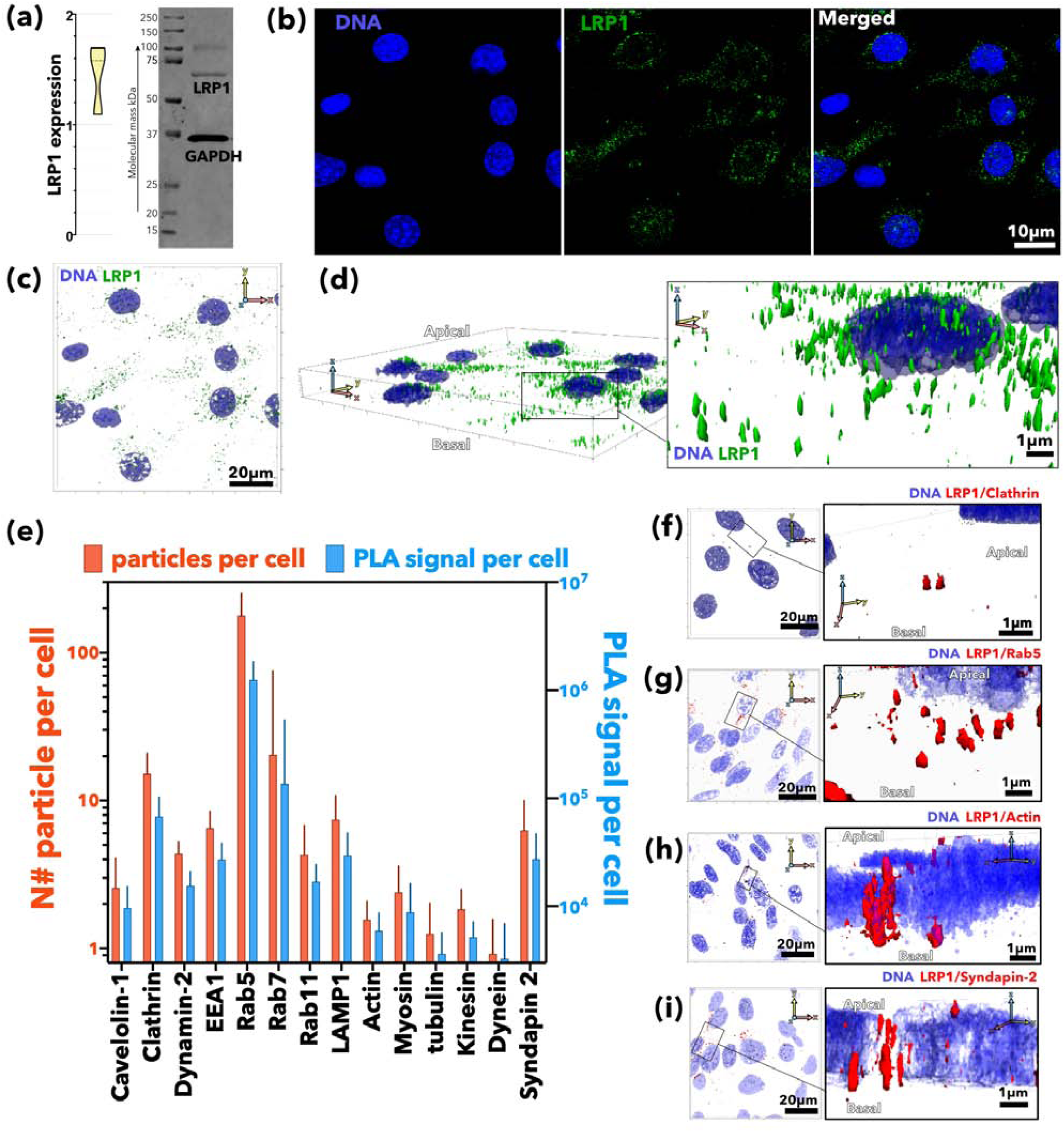
LRP1 intracellular mapping. Expression levels of LRP1 in brain endothelial cells (BECs) assessed by western blot **(a)** and immuno-fluorescence (green) with cell nuclei counter-stained with DAPI (blue) **(b)**. 3D renderings of BECs with both DNA (DAPI in blue) and LRP1 (anti-IgG in green) labelled shown as top view **(c)** and projection **(d)**. Proximity ligation assay (PLA) between LRP1 and several intracellular proteins associated with endocytosis and trafficking reported as number of PLA events per cell and the total PLA signal per cell **(e)**. 3D rendering of BECs showing PLA events between LRP1 and clathrin **(f)**, Rab5 **(g)**, actin **(h)** and syndapin-2 **(i)**.

We next performed a proximity ligation assay (PLA) between LRP1 and several cellular components whose role has been reported during one or more stages of transcytosis [12, 8]. We looked at LRP1 association with early-stage endocytosis effectors (clathrin, caveolin-1, and dynamin-2), main cytoskeleton units (*β*-actin and *α*-tubulin) as well as their corresponding motors (myosin, kinesin and dynein). We also assessed the interaction between LRP1 and early endosomes (Rab5 and EEA1), recycling endosomes (Rab11), late endosomes (Rab7), and lysosomes (LAMP-1). Finally, we also investigated the proximity between LRP1 and syndapin-2, the F-BAR domain-containing protein able to stabilise tubular structures. We performed the assay on polarised BECs imaging them by confocal laser scanning microscopy and collecting about 20 optical sections. The images were then analysed using an *ad hoc* developed algorithm to extract two parameters, the total PLA signal per cell (*PLA*_*C*_) and the number of events (*N* _*E*_) *per* single cell (**Fig.1e**). The *PLA*_*C*_ quantifies whether we have interaction between LRP1 and the targeted protein and estimates the level of such an interaction. The *N*_*E*_, on the other hand, gives us an idea of whether the interaction is distributed across the cell or concentrated in particular loci. Finally, we used the confocal optical sections to generate 3D rendering (**Figs.1f-i**) of the proximity spots to reveal their morphology. The results reported in (**Fig.1e**) show an evident and expected correlation between LRP1 and most endocytic cellular components, but also a particularly strong association with clathrin and Rab5. The 3D rendering showing the proximity spots between LRP1 and clathrin in **Fig.1f** reveals small spots with size at the limit of the confocal resolution, but with morphology suggesting small trafficking vesicles budding out. Similarly, the 3D rendering between LRP1 and Rab5 (**Fig.1g**) resamble endosome morphology and size. We also observed a good association between LRP-1 and *β*-actin, but little or no interaction with *α*-tubulin and all the molecular motors. Interestingly, the LRP1/actin association is concentrated in few events *per* cell and the proximity spots appear as large tubules spanning almost the entire cell thickness (**Fig.1h**). Similar structures accross the cell were also observed between LRP1 and syndapin-2 (**Fig.1i**), which is well known to be associated with tubular membranes [39]. We also confirmed that syndapin-2 is expressed *in vivo* and it is found in several brain cells including BECs (**Fig.2a**). To interrogate tube dimensions and morphology, we then imaged brain sections in super-resolution using stimulated emission depletion (STED) microscopy to have a spatial resolution close to 50 nm. The detailed reconstruction of a single lectin-stained brain capillary is shown in **Fig.2b, c**, and it is evident that syndapin-2 is associated with tubular structures of diameter between 200 to 500 nm and lengths up to a few microns spanning across the endothelial cell (**Fig.2d-f**).

**Figure 2:**
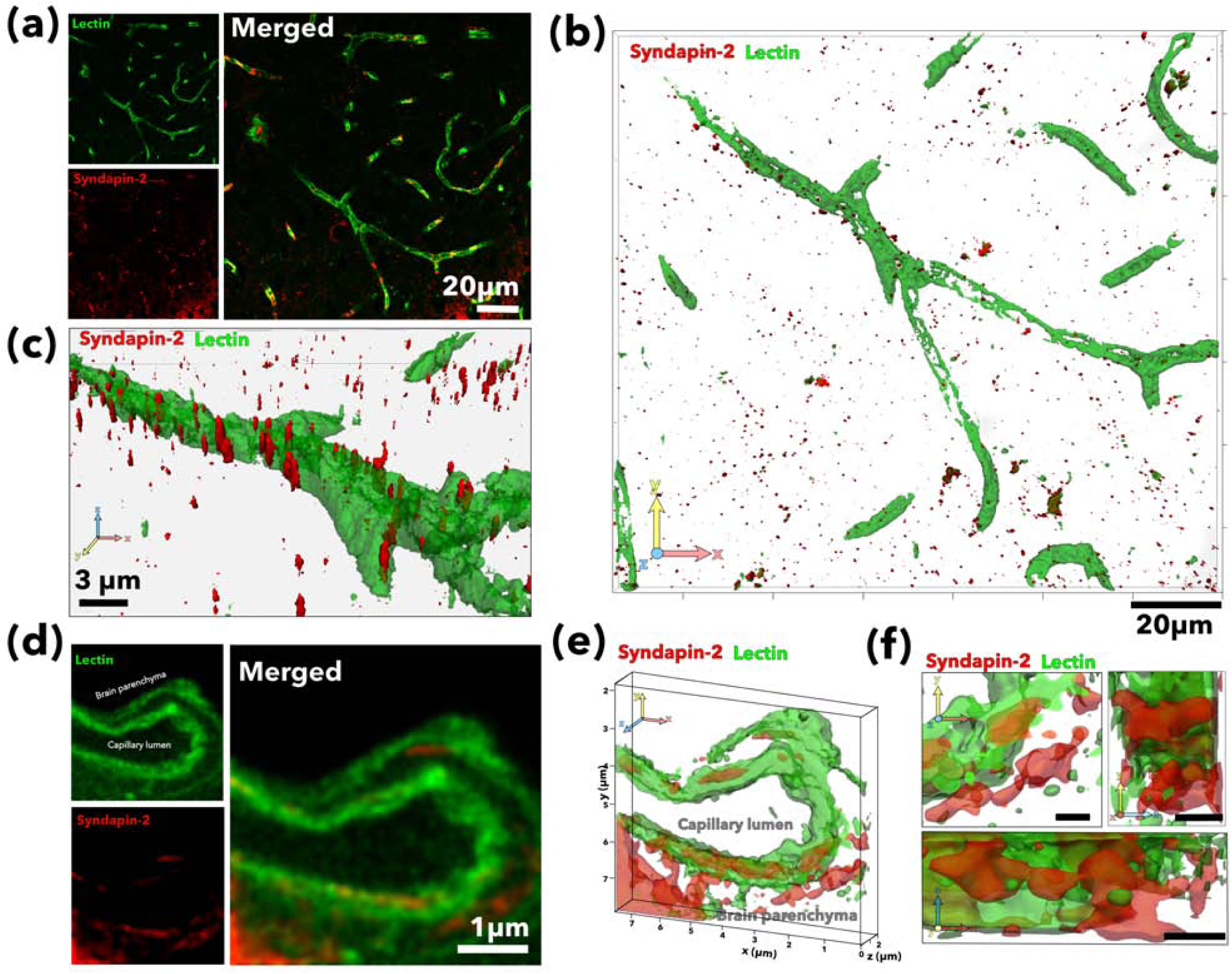
Syndapin-2 expression *in vivo*. Mouse brain cortex histologies imaged by confocal laser scanning microscopy with capillaries stained by lectin (green) and showing expression of syndapin-2 (red) **(a)**. 3D renderings of brain capillary stained by lectin (green) and syndapin-2 (anti-IgG in red) shown as top view **(b)** and close-up projections **(c)**. Section of a single brain capillary stained by lectin (green) and syndapin-2 (anti-IgG in red) imaged by Stimulated Energy Depletion (STED) microscopy with x-y spatial resolution of c.a. 50 nm **(d)** and corresponding 3D rendering showed as projection **(e)** and close-up from top, bottom and side view **(f)**.

### Effect of avidity on BBB crossing

In order to elucidate how the LRP1 moves across the BBB and regulates cargo sorting, we employed one of its most established ligand: angiopep-2. This peptide was derived from the LRP1-binding aprotinin [24] and demonstrated to cross the BBB shuttling anticancer drugs,[40], analgesics [41], RNA[42], DNA [43], and bacteriophage [44]. We demonstrated in both mice [36] and rats [37, 45] that when angiopep-2 is conjugated multivalently to the surface of pH-sensitive POs it augments BBB crossing and enables the intracellular delivery of whole antibodies [36] and neuroprotective peptides [45] into CNS cells. POs are synthetic vesicles formed by the self-assembly of block copolymers in water [46] whose final shape, size and surface topology can be controlled bottom-up [47, 48]. As shown in **Fig.S2**, we produced POs by mixing pristine (**Fig.S2a**) and angiopep-2 modified poly[oligo(ethylene glycol) methacrylate]-(poly (2(diisopropylamino)ethyl methacrylate) (POEGMA-PDPA) (**Fig.S2b**) to make formulations displaying on their surface a different number of ligands and hence with different overall avidities (**Fig.S2c**). The POs referred here as A_*L*_-P, with *L* being the average angiopep-2 ligands number per particle, were produced to be the same size and morphology, as confirmed by both dynamic light scattering (**Figs.S2d-e**) and transmission electron microscopy (TEM) (**Fig.S2f**). We also can infer that within the range of ligand number we used here, POs have almost identical surface chemistry as the highest ligand number *L* = 220, which corresponds to a 12% molar fraction of peptide modified copolymers occupying 6% of the PO external surface area. Effectively, POs become the ideal cargo model to study the shuttling mechanisms across the BBB. To track the POs *in vitro* and *in vivo*, we conjugated cyanine 5 (Cy5) and 7 (Cy7) dyes to POEGMA-PDPA copolymers and mixed them to create the different formulations with a constant concentration of dye. Furthermore, we encapsulated an *ad hoc* synthesised 7-(p-tolyl)-5,6,8,9-tetrahydrodibenzo[c,h]acridine complexed with platinum and dimethyl sulfoxide (PtA2), of which synthesis mechanism and characterisation are shown in **Fig.S3**. This compound was chosen for its superior photostability and metallic nature that allows imaging in super-resolution (STED) and TEM, and also permits for precise quantification in tissues by inductively coupled plasma mass spectrometry (ICP-MS).

We started by measuring the crossing of A_*L*_-P with *L* = 0, 22, 36, 56, 110, and 220 as well as the free angiopep-2, referred here as *L* =1, by administration to the apical compartmental of the *in vitro* BBB model and quantifying the concentration of POs in the basal compartment. Data are collectively shown as a heat map of crossing efficiency (%) as a function of ligand numbers *per* particle and incubation time (**Fig.3a**). It is evident that BBB crossing does not linearly correlate with ligand number, and optimal crossing is shown *L* at = 22 with lower or higher ligand numbers showing a significantly reduced efficiency. From the *in vitro* screening we selected three A*L*-P formulations: *L* = 0, 22 and 110 and the free peptide *L*= 1 for further testing in mice. After 2 hours of intravenous injection, we perfused the animals with saline solution to remove the excess of blood, harvested the whole brains and imaged them using the *in vivo* imaging system (IVIS) (**Fig.3b**). The same brains were subsequently processed to extract the parenchymal fraction and then quantify the percentage of injected dose (% ID) of A*L*-P present in the tissue using near-infrared fluorescence spectroscopy (**Fig.3c**). Both methods show that while all angiopep-2 formulations enter the brain and can be found at relatively high concentrations across the BBB, the formulation with the most effective crossing is again A_22_-P, in agreement with the *in vitro* data. Such a non-linear dependence of the ligand binding energy on BBB crossing rate was also reported for the targeting of the transferrin receptor [49, 50, 51] and glucose transporter-1 (GLUT1) [52].

**Figure 3:**
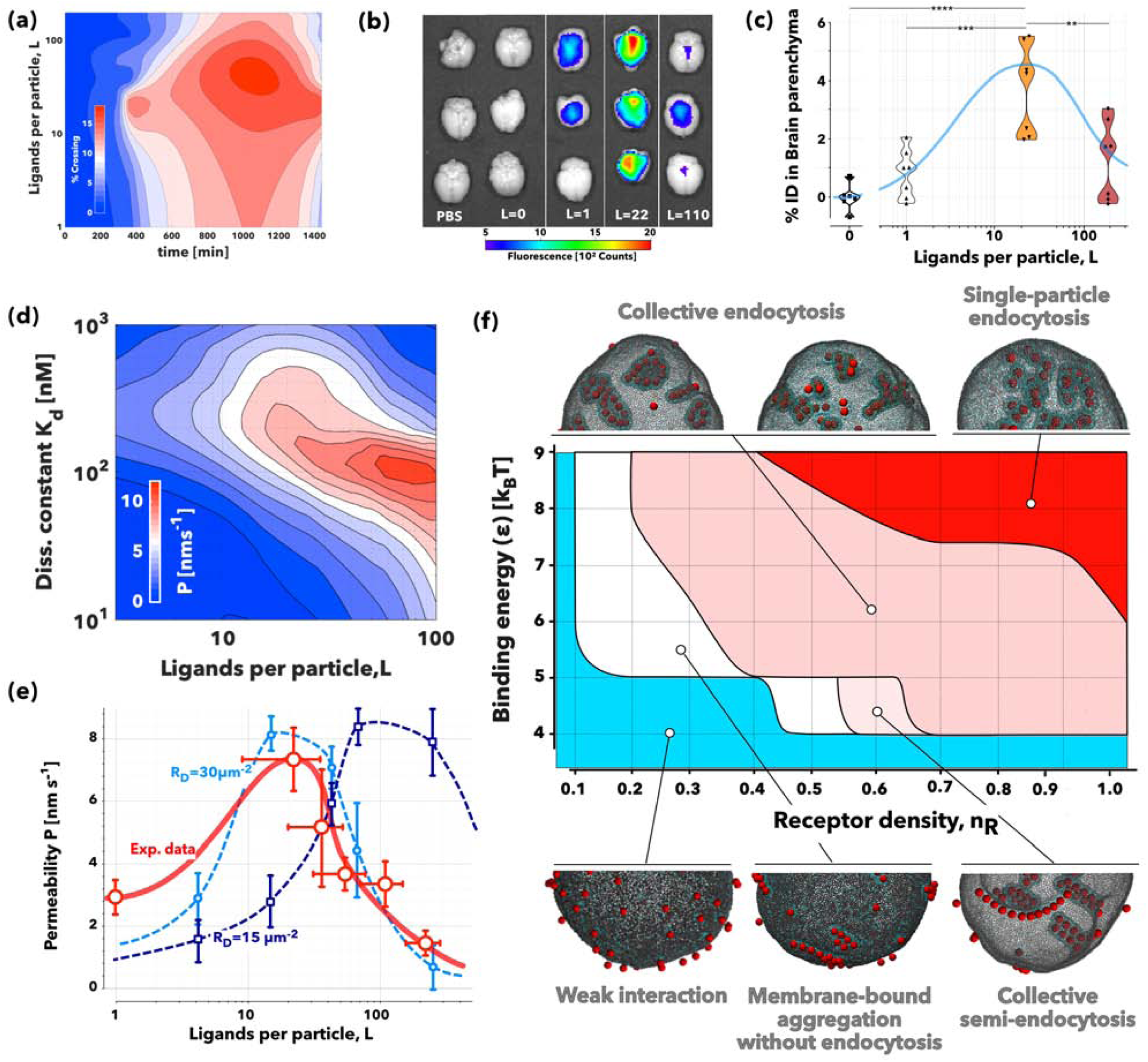
Ligand avidity versus BBB crossing. Heat-map showing the experimental measurement of % of A_*L*_-P (POs) crossing as a function of incubation time and ligand number per particles (*L*) **(a)**. *Ex vivo* fluorescent photographs of whole murine brains imaged 2 hours after intravenous injection of PBS, pristine POs (*L=0)*, free angiopep-2 peptide (*L*=1), A_22_-P or A_110_-P **(b)**. Violin plots showing the quantification in the brain parenchyma of the various preparations tested. ** *P* < 0.01, *** *P* < 0.001, **** *P* < 0.0001. One-Way ANOVA with (*n* = 6). Concentration of angiopep-2 functionalised cargo expressed as % injected dose per gram of tissue as a function of the number of ligands **(c)**. Heat map of the apparent permeability, *P*, obtained from agent-based simulations as a function of the ligand number per particles and the single ligand dissociation constant, *K*_*d*_ with the LRP1 receptor **(d)**. Comparison between apparent permeability, *P*, across BBB experimental data (red markers and solid line) and simulation (blue markers and dashed lines) calculated for two different receptor densities and single ligand dissociation constant, *K*_*d*_ = 300 nM **(e)**. Note the control pristine POs apparent permeability was subtracted to the other formulations to remove passive diffusion. Phase diagram showing different regimes of nanoparticle aggregation across the receptor densities and nanoparticles-receptor affinities expressed in *k*_*B*_*T* (with *k*_*B*_ being the Boltzmann constant and *T* the temperature) as observed in molecular dynamics simulations **(f)**.

From a theoretical standpoint transcytosis involves five major stages: binding, endocytosis, trafficking, exocytosis and unbinding. Efficient transcytosis requires the formation of ligand/receptor bonds that last enough for it to be trafficked across, yet the higher the ligand binding energy, the lower is its ability to detach once across to the other side. Therefore, a balance is required to form and maintain sufficiently strong bonds to enable binding and endocytosis, but also a sufficiently weak bond to allow unbinding and release. Such an approximation allows creating an *in silico* model to stimulate transcytosis, see Supporting Information. We employed Flexible Large-scale Agent-based Modelling Environment (FLAME), a generalised agent-based modelling plattform, that models the behaviour of individual POs undergoing Brownian motion, binding to endothelial cells, crossing the cells by transcytosis and being released into the basal compartment [53]. We designed the model based on the geometry of the transwell insert used in the *in vitro* experiments (**Fig.S1a**), and the BECs were modelled as a uniform 2-*μ* m thick layer at the top of the insert. We also modelled POs with different ligand numbers and different individual ligand-receptor dissociation constants, starting them at time zero in the aqueous apical phase. POs were subjected to Brownian motion and bound to cells according to the multivalent-avidity binding model described in Supporting Information. The particles were allowed to go through the different stages of transcytosis as described in **Fig.S4a** and the number of POs that crossed the BECs was measured. We thus used **Equation 1** to calculate the apparent permeability and plotted it as a function of both ligand number *per* particle (*L*) and the single ligand/receptor dissociation constant (*K*_*d*_). We employed models for nanoparticles with radius *R* = 20 and 50 nm, as well as receptor densities *R*_*D*_ = 15 and 30 *μm* ^−2^, respectively. According to the simulations, there is a non-linear dependendence between ligand number and binding strength (**Fig.S4b**), whereby the optimal transcytosis is obtained in a “Goldilocks” regime of avidity, *i*.*e*. not-too-strong and not-too-weak, and it is independent of the particle size or receptor density. We selected both size and receptor density to match our *in vitro* experimental data, and within such a range, our simulations suggest that bigger particles and larger receptor density lead to improved transcytosis. In **Fig.3d** we plot the apparent permeability across the BBB as a function of ligand numbers *per* particles (*L*) and the dissociation constant of the ligand/receptor binding (*K*_*d*_) for particles with radius *R* = 50 nm and receptor density *R*_*D*_ =30 *μm*^-2^, that is very close to what we recently estimated using the super-selective theory [54]. We know from previous work that angiopep-2 has a dissociation constant *K*_*d*_ = 313nM [55], and using this we can thus compare the simulations at similar dissociation constant with the experimental data. In **Fig.3e**, we plot the experimental apparent permeability (in red), measured from the data in **Fig. 3a** and the simulations with *K*_*d*_ = 300 nM, particle with size *R* = 50 nm, and receptor densities *R*_*D*_ = 30 *μm*^-2^ or *R*_*D*_ = 15 *μm*^-2^, respectively. The experimental and simulation data show broad agreement, and indeed the Goldilock effect is reproduced experimentally at similar values to those we observed computationally.

Finally, we complemented both computational and experimental permeability measurements by performing molecular dynamics (MD) simulations to capture the effect of avidity on membrane topological changes and nanoparticle aggregation dynamics. We used a well established coarse-grained membrane surface patch (**Fig.S5.a**) on an equilibrated spherical membrane and varied the receptor density and the nanoparticle to membrane binding energy (*ϵ*) expressed in *K*_*B*_*T* with *K*_*B*_ being the Boltzmann constant and *T* the temperature. The latter represents the depth of the potential well in the attractive interaction between nanoparticles and ‘receptor’ membrane beads (see **Equation 3** in Supporting Information). A different initial nanoparticle distribution was randomly chosen for each simulation and each parameter pair used the same set of 6 different initial nanoparticle distributions. The receptor density was represented in the model by the ratio of ‘receptor’ membrane beads to the total number of membrane beads. The simulation results are summarised in **Fig.3f**, and it is evident that across all receptor densities, no clear binding of the nanoparticles was observed for low binding energy. Some ‘receptor’ beads clustered around individual adsorbed nanoparticles but the nanoparticle-receptor adhesion was too weak to drive any interaction. As the binding energy increases, the nanoparticles bind to the membrane. While their relative adhesion energy is converted into membrane deformation, this is not sufficiently strong to induce full endocytosis. Nonetheless, progressively more particles bind the associated membrane deformations forming linear aggregates. The anisotropic aggregation is the consequence of the trade-off between nanoparticle-receptor adhesion and the membrane’s resistance to deformation [56]. Higher binding energy results in the linear aggregates being internalised as tubular aggregates. These can co-exist on a membrane together with membrane-bound tubular aggregates and internalised tubular aggregates. At higher receptor densities, lower binding energies are required for the nanoparticles to form tubular and linear aggregates. However, at high receptor density and high binding energy, the particles have sufficient adhesion to create singular deformation and enter *via* descrete endocytic events. The pseudo-phase diagram in **Fig.3f** shows the limits of the different regimes observed in the simulations, and the collective processes leading to different outcomes. As shown in **Fig.S5b**, tubulation results in a high number of cargo units transported *per* single event, while higher binding energy and receptor density correspond to fewer number of particles *per* internalised carrier. Interestingly, the latter process is more efficient in internalising the nanoparticles reaching almost 100%, while the collective process that occurs at lower binding energy and receptor density achieves only a much lower percentage of internalisation (**Fig.S5c**). All together the MD simulations add another dimension to the avidity effect showing that different binding energies drive alternative membrane deformations, and that these lead to a different endocytic initiation.

The data in **Fig.1** showed that LRP1 is associated with several endocytic and trafficking elements, suggesting that the receptor is physiologically processed in endosomes and lysosomes, but it is also shuttled in tubular structures stabilised by syndapin-2. We thus repeated the PLA assay between LRP1 and the same proteins only this time BECs were exposed to angiopep-2 peptide (*L* = 1) and A_*L*_-P formulations, *L* = 22 and *L* = 110. The data reported in **Fig.4a** as a variation between the treated and untreated cells reveal how the avidity of the ligand for LRP1 affecs the localisation of the receptor within the cells. At an early incubation time (0.25 hour), the single peptide (*L* = 1) reduces the proximity events between LRP1 and clathrin by more than 5 times, while prolonged incubation promotes the association of the receptor with the late endosome marker Rab7. The two A_*L*_-P formulations have a more dramatic effect. Incubation with *L* = 22 prevents the interaction of LRP1 with all the endo-lysosomal compartments at any time point, with Rab5 showing the most notable decrease. Interestingly, the presence of *L* = 22 at 0.25 hour also increases the interaction of the receptor with actin, tubulin and clathrin. On the other hand, incubation with *L* = 110 increases LRP1 interaction with both Rab7 and Rab11 in the short-term, while it constantly reduces the association with syndapin-2. At later time points, incubation with *L* = 110 incubation with also decreases the proximity between LRP1 and tubulin or clathrin. Note that oscilations of the values between −5 and +5 were considered as physiological fluctuations. Our data suggest two trends; one is the association of LRP1 with syndapin-2 and one is with Rab5. We then plotted the ratio of the relative interactions between LRP1/Rab5 and LRP1/syndapin-2 (Rab5/syndapin-2) as a function of incubation time and ligand number (**Fig.4b**). While angiopep-2 peptide does not alter the Rab5/syndapin-2 ratio, both *L* = 22 and *L* = 110 do but with opposite trends. The *L* = 22 pushes the interaction of LRP1 toward syndapin-2 for all time points, while *L* = 110 biases LPR1 toward the endosomal protein Rab5. As we expected that LRP1 association with endo-lysosomal markers should result in its degradation, we assessed its levels of expression over time following incubation with *L*=1, *L*=22, and *L*=110 (**Fig.4c**). The WB results show that LRP1 is unaltered after up to 1 hour incubation with angiopep-2 and *L* = 22. Contrarily, exposure to *L* = 110 results in a fast reduction of LRP1 expression, which then recovers to physological levels after 2 hour of incubation. Most interestingly, we observed a 2-fold increase in LRP1 expression after 2 hours incubation with *L* = 22.

**Figure 4:**
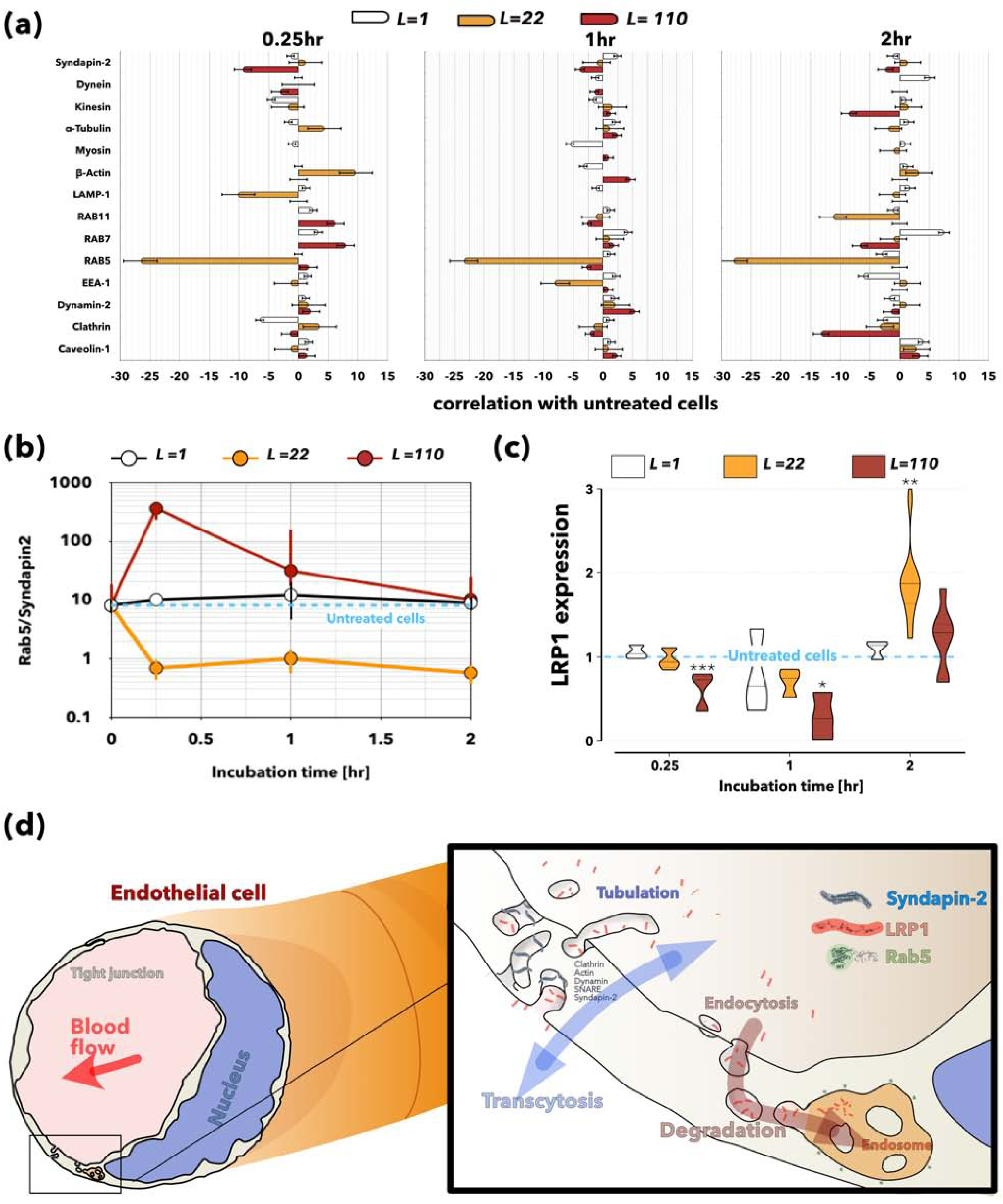
LRP1 subcellular localisation and expression as a function of avidity. Deviation of the number of proximity events measured by a proximity ligation assay (PLA) between untreated endothelial cells and treated for 0.25, 1 and 2 hours incubation with free angiopep-2 peptide, *L* = 1, and A_*L*_-P with *L* = 22 and *L* = 110 **(a)**. Note that zero corresponds to no variation, while positive and negative values indicate up- and down-regulation, respectively. Ratio between LRP1/Rab5 and LRP1/Syndapin-2 number of proximity events for the different treatments with free angiopep-2 peptide, *L* = 1, and A_*L*_-P with *L* = 22 and *L* = 110 with Rab5/Syndapin2 being 10 for the untreated cells **(b)**. Western blot measuring the LRP1 expression relative to the untreated cells for free angiopep-2 peptide, *L* = 1, and A_*L*_-P with *L* = 22 and *L* = 110 measured at different incubation times with 0.25, 1 and 2 hours. * *P* < 0.05, ** *P* < 0.01, *** *P* < 0.001. One-Way ANOVA with (*n* = 6) **(c)**. Note that LRP1 expression is normalised to the loading control. Diagram showing the syndapin-2 mediated transcellular route and the intracellular degradation of LRP1 **(d)**.

Overall, both PLA and WB analysis suggest that LRP1 can follow two different intracellular pathways across BECs and their schematics are shown in **Fig.4d**. One pathway is mediated by syndapin-2, *β*-actin and possibly clathrin. Here, LRP1 shuttles across tubular carriers from apical to basal and *vice-versa* avoiding endolysosomal degradation and sorting. The other pathway is a conventional endocytosis where LRP1 enters the cells and gets trafficked to endosomes and lysosomes where it is degraded. We also show that BECs favour one pathway over the other depending on the cargo avidity, with intermediate ligand numbers pushing more to the syndapin-2 pathway associated with tubular deformations, while the higher number of ligands and avidity pushes the cargo more towards endosomal sorting. The result, as demonstrated above, is that in the former the cargo is shuttled across efficiently, while in the latter it is degraded and possibly consumed by the same BEC. In other words, using the same receptor but controlling its clustering and association, BECs can discriminate between nutrients destined to the brain or to themselves.

### Transcytosis mechanism

To further shed light on the novel shuttling mechanism, we employed a combination of qualitative and quantitative confocal microscopy in conjunction with antibodies and small molecule inhibitors against proteins of interest. First, we confirmed that Cy5-labelled A_22_-P interact with the LRP1 in BECs by performing immuno-fluorescence of LRP1 after incubation with POs. In **Fig.S6a**, confocal micrograph shows the association of A_22_-P and LRP1 after 60 minutes, and with the PO signal being detected in different areas of the transwell as cytoplasmic puncta. At a higher magnification, many puncta of LRP1 and A_22_-P are visibly colocalising with the cell membrane (**Fig.S6a1**). To elucidate the extent and time that A_22_-P associates with LRP1, colocalisation was quantified with Pearson’s correlation coefficient, *r*. After 10 minutes of incubation, *r* = 0.78 for A_22_-P and LRP1 with colocalisation remaining high as 0.78 after 120 minutes (**Fig.S6a**). The continuously high *r* values indicate that A_22_-P remains associated with the intracellular domain of LRP1 all along the intracellular stages of transcytosis. To further confirm this, we co-incubated the A_22_-P with the free peptide to provide an insight on whether transcytosis is more efficient when angiopep-2 is alone or when attached to the POs. Quantification of fluorescence is shown in **Fig.S6b**. After 10 minutes of co-incubation, A_22_-P fluorescence is of similar intensity of angiopep-2 and much lower than that of A_22_-P after 10 minutes with no competing ligand. Free peptide fluorescence remains similar to levels without competition. Such results show that angiopep-2 and A_22_-P compete for LRP1 binding and endocytosis, as expected, but also that the free peptide inhibits POs internalisation more than *vice versa*. When co-incubated, the intensity of A_22_-P and angiopep-2 are both markedly higher at 60 minutes compared to when added without competition. However, competition for A_22_-P shows a biphasic shift in behaviour compared to the A_22_-P only control: decreased endocytosis at 10 minutes and increased intracellular residence, *i*.*e*. decreased exocytosis at 60 minutes. The biased inhibition of A_22_-P transcytosis rather than angiopep-2 may be due to more rapid or efficient endocytosis, intracellular trafficking and exocytosis pathway occurring for A_22_-P than for angiopep-2.

We subsequently studied the mechansims of endo- and exocytosis of Cy5-labelled A_22_-P during transcytosis. Confocal studies suggested that clathrin, but not caveolin, is involved in the mechanism of internalisation of A*L*_22_-P (**Fig.S6c,d**). High manification confocal images in **Fig.S6c** demonstrate that A*L*_22_-P fluorescence is closely associated with clathrin after 60 minutes of incubation. However, this data is qualitative and thus only an indication that clathrin is involved in transcytosis of A*L*_22_-P. We performed similar experiments to evaluate the association of Cy5-labelled A*L*_22_-P with caveolin-1, and as shown in **Fig.S6d**, a partial overlap was observed initially at 10 minutes of incubation. However, 3D z-stack projections in **Fig.S6e** display no apparent colocalisation at 10 minutes. A few cytoplasmic puncta with fluorescence overlap were observed at 60 minutes. However, *r* values for A_22_-P and caveolin-1 remained low along the time with *r* = 0.2 and −0.02 at 10 and 60 minutes, respectively. Overall, these findings fail to show a role for caveolae as essential structures for apical and basal transcytosis, particularly, as a higher colocalisation would be antecipated at 10 minutes when the majority of transcytosis is occuring. Cytoskeletal motor proteins can quickly transport cargo from one side of a cell to another and were therefore of particular interest for their potential involvement in transcytosis. We thus investigated the role of actin in BEC transcytosis by colocalisation of Cy5-labelled A_22_-P with phalloid-488 (an established marker for F-actin). Confocal images are displayed in **Fig.S6f**, with a magnification of an area of interest (**Fig.S6f1**), along with *r* values at 10, 30 and 60 minutes for A_22_-P and F-actin. The data suggest that actin has a role in transporting POs from the apical to basal membrane within the first few minutes of endocytosis. The timescale of BEC transcytosis and unconventional intracellular trafficking pathways prompted us to further explore the identity of intracellular transport vesicles as well as membrane deformation mediators in transcytosis. Small molecule inhibitors of endocytosis or exocytosis were used in conjunction with live-cell imaging to obtain transwell z-stacks. Incubation with dynasore, a cell-permeable inhibitor of dynamin, impaired transcytosis and caused Cy5-labelled A_22_-P to remain stuck on the BEC surface (**Fig.S7a,b**). These effects were reversible upon removal of the inhibitor as the A_22_-P were visible both inside cells and in transwell membrane pores. Dynamin may, therefore, be a required cellular component of the internalisation stage of transcytosis in BECs. In a separate experiment, N-ethylmaleimide (NEM) was used to inhibit NEM soluble factor (NSF) in order to inhibit exocytosis indirectly. A_22_-P remained aggregated on top of the cells after incubation for 60 minutes (**Fig.S7a,b**). Thus, NSF appears to be a central component not only for exocytosis of cargos once inside the cell but also for endocytosis. Washing out NSF provided no recovery in POs transcytosis (data not shown). To further explore the role of NSF and SNAREs in transcytosis, a cell membrane cholesterol depletion method was used to disrupt lipid raft containing SNAREs. Cells were pre-incubated for 60 minutes with methyl-*β*-cyclodextrin (CD) added to either the apical or basal compartment of the transwell. A cholesterol quantification assay revealed a slight asymmetry in measured free cholesterol in the media in the apical and basal compartments (**Fig.S8a**). Depletion of cholesterol in the apical or basal membrane resulted in an approximately 2-fold or 3-4-fold increase in cholesterol released into the apical or basal side of the transwell, respectively (**Fig.S8b**). Such an effect may be indicative of a stronger effect of cholesterol depletion on the basal membrane. Confocal images were acquired of Cy5-labelled A_22_-P incubated for 60 minutes in BECs with CD added to the apical or basal side of the transwell (**Fig.S8c**). Basal membrane cholesterol depletion showed an increase in intracellular A_22_-P after 60 minutes compared to untreated cells, which may be due to the ability of cells to do endocytose but not exocytose the cargo. Altogether, these findings implicate the involvement of dynamin and also NSF in the LRP1-mediated cargo internalisation stage of BECs transcytosis. Depletion of cholesterol in the basal side of BECs inhibited exocytosis but not endocytosis demonstrating the essential role of cholesterol in transcytosis. We next assessed whether the trafficking from apical to basal involves sorting into endosomes and acidification, as we already demonstrated [36], A_22_-P do not colocalise with endosome and lysosomes crossing the BECs without losing integrity. Here, we represent confocal images acquired of A_22_-P in BECs fixed and stained for Rab GTPases of endosomal organelles in **Fig.S9a**. Surprinsigly, there was no co-localisation between POs and any of the markers at any time investigated. Colocalisation quantification (**Fig.S9b**) indicated no association between A_22_-P and Rab5, Rab7, Rab11 and LAMP-1. On the contrary, *r* values displayed a negative trend implicating negative association, *i.e*. exclusion of A_22_-P from these organelles.

Finally, we confirmed the colocalisation between A_22_-P and syndapin-2 in our *in vitro* BBB model. In **Fig.5a**, 3D rendering of polarised BECs imaged 30 minutes after incubation with the Cy5-labelled A_22_-P (red) shows very effectively that A_22_-P cross the cell forming tubular structures, and that these are coated with syndapin-2 (in green). To further show the involvement of syndapin-2 on the transcytosis of A_22_-P, we modulated the expression of syndapin-2 on BECs and assesses the transport of A_22_-P across an *in vitro* BBB model (**Fig.S10**). Specifically, we performed short hairpin RNA (shRNA) on bEnd3 to knockdown syndapin-2 generating a stable cell line expressing significantly less syndapin-2, as confirmed by WB (**Fig.S10a**). When cultured onto collagen-coated transwells, these syndapin-2 knockdown bEnd3 showed permeability *P*_4*kDa*_ = 25.6 and *P*_70*kDa*_ = 5.4 nm s^−1^, which are similar to the values obtained for bEnd3 transfected with a control shRNA (**Fig.S10b**). We then assessed the transport of A_22_-P across BECs expressing different levels of syndapin-2. In **Fig.S10c**, we clearly observe a 2-fold decrease in the apparent permeability of A_22_-P from apical to basal side when compared to bEnd3 expressing normal endogenous levels of syndapin-2. These results further indicate the involvement of syndapin-2 in the transport across BECs. We complemented the colocalization of syndapin-2 and A_22_-P with animal studies where we injected either A_22_-P or pristine POs loaded with PtA2. In **Fig.5b**, the *ex vivo* fluorescent photographs of whole brains extracted from healthy mice 30 minutes post-injection show the effective delivery of the dye by functionalised POs. PtA2 has unique fluorescence characteristics with a wide Stoke shift and etremely bright emission allowing us to visualise the POs penetration with high sensitivity. Most importantly, the metallic nature of the dye allows quantification of its biodistribution by ICP-MS. The graph in **Fig.5c** shows an extremely effective delivery of the dye into the brain with a staggering ratio brain/liver of about 8.2 opposite to the pristine POs where the majority of the dye is found in liver and spleen. Such a high concentration of dye allow us to visualise A_22_-P penetration in the brain capillary by TEM (**Fig.S11**) and STED. The histology in **Fig.5d** demonstrates that A_22_-P cross the brain endothelium (stained with lectin in green) *via* the formation of tubules as shown in the region of interest 1, 2 and 3. We then imaged brain sections collecting 30 optical slides, and the corresponding 3D renderings are shown in **Fig.5e** where the A_22_-P loaded with PtA2 (red) are imaged alongside the capillary walls (green) and syndapin-2 (blue) with improved spatial resolution. The rendering show very well that A_22_-P colocalise into tubular structures coated by the F-Bar protein syndapin-2 with dimensions in the aggrement of what we observed *in vitro* and by the simulations.

**Figure 5:**
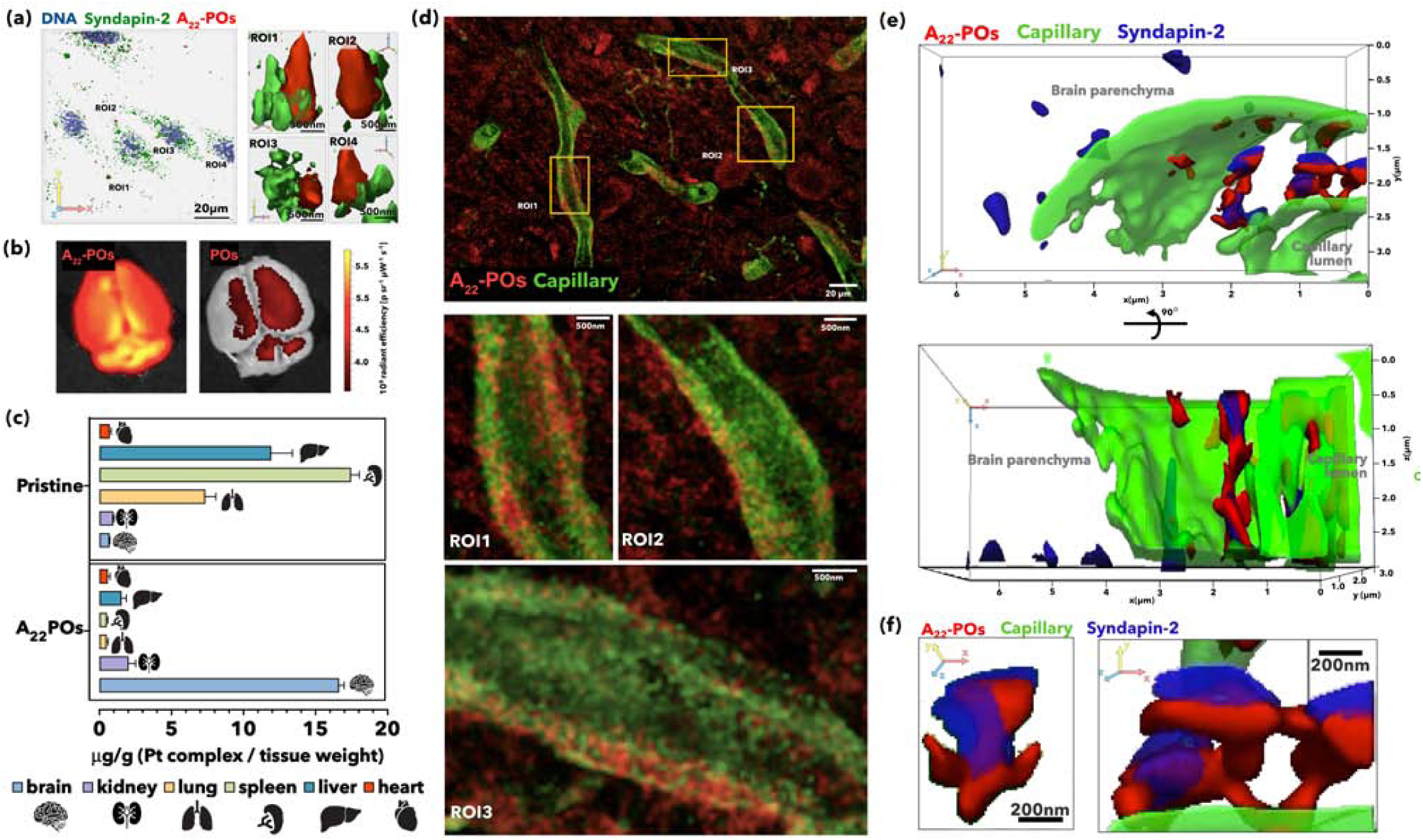
Syndapin-2 mediated transport. 3D rendering of confocal laser scanning micrographs of polarised brain endothelial cells (BECs) incubated with A_22_-P (red). Cell nuclei were stained with DNA binding DAPI (blue) and syndapin-2 is showing in green (anti-IgG) **(a)**. Fluorescence photograph of *ex vivo* whole mouse brains imaged 30 minutes after intravenous injection of A_22_-P and pristine POs loaded with PtA2 **(b)**. Pt tissue concentration in brain, kidney, lung, spleen, liver, and heart expressed as *μ*g/g and measured by inductively coupled plasma mass spectrometry (ICP-MS). Tissues were collected 30 minutes after intravenous injection of A_22_-P and pristine POs both loaded with PtA2 **(c)**. Stimulated Energy Depletion (STED) micrographs of coronal brain sections showing the distribution of PtA2 loaded A_22_-P (red) 30min post-injection with capillary stained by lectin (green). Three different regions of interests (ROIs) show the detail of the tubulation across the BECs **(e)**. 3D renderings shown as projections of STED micrographs of brain capillary (lectin in green) showing the detail of PtA2 loaded A_22_-P (red) and syndapin-2 (anti-IgG stained in blue) **(f)**. Details of the tubule formed by the PtA2 loaded A_22_-P (red) surrounded by syndapin-2 (anti-IgG stained in blue) **(g)**.

### Transcytosis dynamics

As described above, A_22_-P formulation is extremely effective in crossing the BBB and thus makes the ideal tool to study the transcytosis dynamics. In **Fig.S12a**, we show optically reconstructed sections of fixed polarised BECs incubated at different times with A_22_-P and stained for claudin-5. A_22_-P interact very quickly with the cells, and within 10 minutes, we observe POs crossing into the porous membrane. Interestingly, as shown by the co-staining with claudin-5, A_22_-P seem to concentrate throught the cells, and almost no fluorescence is observed in the tight junctions indicating that A_22_-P diffusion across endothelial cells is transcellular rather than paracellular. We then measured the average fluorescence across apical to basal direction at different time points and disclosed that the transport dynamics are fast and occur within the first hour (**Fig.S12b**). We thus stained polarised BECs with both DNA (DAPI) and membrane stain (CellMask*™*) and imaged the binding and crossing of A_22_-P using real-time 4D (xyzt) live-cell imaging. At first, we collected 3D sections every 8.2 seconds to generate **Video S1**. In **Fig.6a**, three sequential 3D renderings show that A_22_-P (red) interact with the cell membrane forming several clusters that rapidly evolve into tubular endocytic events. Over the duration of the video (40 minutes) we counted a total of 250 complete events, which are shown in the 3D rendering in **Fig.6b** colour coded according to their occurrence. The time sampled rendering shows that are no events over the nuclear and perinuclear endoplasmic reticulum, and most of them are stochastically distributed over the remaining cell surface with a considerable level of overlapping of events occurring at different times but in the same spot. A_22_-P fluorescence intensity and the number of events were analysed as a function of time and zeta averaged across the full cell tickness (**Fig.6c**). Furthermore, each event was analysed by measuring its radius *r*_*e*_ and height *h*_*e*_ as a function of time (**Fig.6d**), and one examplary event is shown in its 3D renderings in (**Fig.6e**). Early stages being with few puncta visible on the cell surface (**Fig.6e1**) with size below the capability of confocal laser scanning microscopy. The puncta assemble forming clusters with areas ranging from a few to 70 *μ*m_2_, with an average of about 25 *μ*m2. The lateral growth of the clusters stops while they mature into tubular structures with a height up to 6 *μ*m, and an average radius of 2.52 *μ*m. After reaching an average aspect ratio of 1.8, the tubules disappear from the field of view faster than our time-resolution allowing us to partially capturing the crossing. To capture the crossing dynamics, we also run experiments at faster acquisition times where each optical section is collected in 2 seconds. The **Video S2** and the snapshots in **Fig.6f** show a strong interaction between A_22_-P and the BECs again. While the imaging quality is inevitably compromised, making the visualisation of small early-stage events challenging, we can capture the dynamics of large tubular structures moving from one side to the other. We can thus measure the time each tubule takes to go across, defined here as crossing time *τ*_*crossing*._. In **Fig.6g**, we plot the normalised mean square displacement (MSD) averaged across 51 events as a function of the normalised time calculated as *τ\** = *t*/*τ*_*crossing*._. The data show that the tubule MSD follows a two-regime trend; at early stages it is linear with time indicative of diffusional processes, while at later stages the trends become parabolic typical of a ballistic process. The two dynamic analysis together allows us to identify four stages of transcytosis: (i) clustering, (ii) tubulation, (iii) fission, and (iv) crossing. At early stages, the cargo is sorted and clustered in an average of 13.5 seconds, where clustering time (*τ*_*clustering*_) was measured when *r*_*e*_ ≥ *h*_*e*_ (**Figs.6h**). Then, the tubulation starts and it occurs in between 20 and 160 seconds, where *τ*_*tubulation*_ is measured when *r*_*e*_ < *h*_*e*_ (**Figs.6i**). This time variance is the reflection of two kinetic processes that we observed. In some cases, the arrest of lateral growth of the clusters leads to an immediate tubulation, while in other instances the tubules roam on the cellular surface for several seconds before disappearing. Next, the tubes fission moving from one side to the other of the cell with a mean value of 110 seconds, where fission time (*τ*_*fission*_) is calculated as *τ*_*clustering*_ +*τ*_*tubulation*_ (**Figs.6j**). The crossing time defined as above is less spread with a mean value of about 15 seconds (**Figs.6k**).

**Figure 6:**
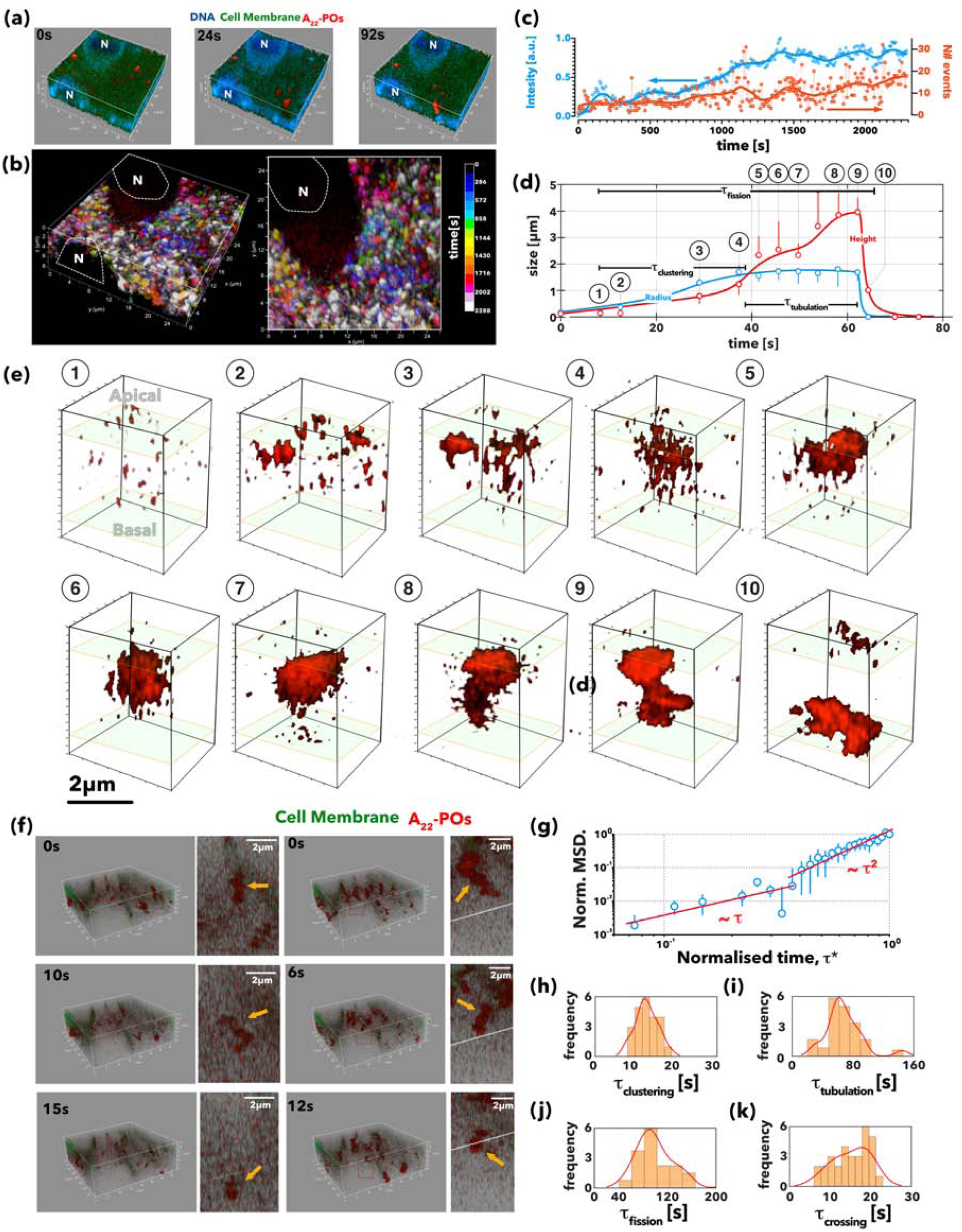
4D microscopy of transcytosis. 3D rendering at three different times extracted from 4D (xyzt) live imaging of brain endothelial cells (BECs) stained for cellular membrane CellMask*™* (green) and nuclei (blue) and incubated with Cy5-labelled A_22_-P (red) **(a)**. 3D renderings of the same cell with each event colour coded by its occurrence within periods of 144 seconds **(b)**. Graphs showing the red channel fluorescence intensity and the number of events (threshold in the red channel) as a function of time and zeta averaged across the full cell thickness **(c)**. Each event radius and length is monitored over time the average values across 20 events are plotted as a function of time **(d)**. The corresponding 3D renderings of the single events show an evolution from few few puncta to large clusters, to membrane bound tubulations, to tubular carriers **(e)**. Two sequences of 3D renderings extracted by fast 4D videos of the tubular carriers filled up with Cy5-labelled A_22_-P (red) crossing from one side to other BECs, note the cell membrane is stained by CellMask*™* (green) **(f)**. Normalised mean square displacement as a function of normalised time *τ*\* = *t*/*τ*_*crossing*._ where *τ*_*crossing*._ is the time each event takes to fully cross from apical to basal and vice-versa **(g)**. Distribution of *τ*_*clustering*_ **(h)**, *τ*_*tubulation*_ **(i)**, and *τ*_*fission*_ **(j)** measured from the graph in **(d)** and *τ*_*crossing*._ **(k)** measured from the graph in **(g)**.

We repeated the live-cell imaging of polarised BECs using, this time, STED microscopy and A_22_-P encapsulating the PtA2 to achieve a spatial resolution close to 40 nm. BECs incubated with PtA2-A_22_-P were monitored in STED mode every 6.7 seconds for about 6 minutes. In **Fig.7a** and **Videos S3, S4**, and **S5**, we show the corresponding 3D renderings colour-coded in terms of depth. We also plotted the z-averaged fluorescence as a function of time showing a remarkable periodicity in the formation of transcytotic events with a similar period to those we observed in **Figs.6c**. STED resolution allow resolving the single nanoparticles, as shown in detail in **Fig.7b**. The most striking revelation is that the transcytotic event emerges as an assembly of many tubular units each having an average diameter of about 100 ± 20 nm and a length varying from few hundreds of nanometres to few micrometres. In **Fig.7c**, we show the evolution of these tubes from a few nanoparticles wide to large interconnected networks. An interesting observation is that the events appear very symmetrical, starting from both apical and basal sides, and growing until they are connected *via* a network of discontinuous tubes. It is important to point out that in both conventional and super-resolution imaging, we observed a dissociative tubule formation with fission preceeding fusion and hence we did not observe a single tubule spanning from apical to basal. As we show in **Fig.3f**, the intermediate binding energy results in nanoparticles forming tubular aggregations, which are then internalised into the membrane (**Fig.7d**). Typically, the nanoparticles first formed short linear surface aggregates that acted as nucleation seeds and grew in length. At some point, the linear aggregate buckles into the membrane, forming a membrane-bound tubular aggregation wrapped in an envelope of ‘receptor’ beads that protruded into the inside of the membrane. The membrane-bound aggregations would then be internalised, forming a separate vesicle inside the membrane. **Figs.7d** shows that once the membrane buckles and starts deforming, it acts as a sink for the other nanoparticles bound to the membrane resulting in the formation a relatively long tubule. In most cases, the tubes undergo fission and severe from the membrane, as shown in **Figs.7e**. The membrane “sinks” can attract more than one linear membrane-bound aggregate resulting in the formation of tubules that are two or more nanoparticles thick **Figs.7f**. Immediately after their formation, but preceding any further growth or interactions, the membrane-bound tubular aggregates could be grouped into three different morphologies (**Figs.7g**) depending on whether they are 1, 2 or 3 nanoparticles wide. Variations in the exact structure of the membrane-bound tubular aggregates were often seen, with aggregates of widths above 3 nanoparticles occasionally observed to arise from the merging of smaller aggregates. On one occasion, with an increased number of nanoparticles (200 instead of the usual 105), we observed a particularly large tubular structure (**Figs.7d**). It was speculated that on a larger computer model, more aggregates could cooperatively take part in tubular growth interactions to give rise to even larger tubular aggregations of nanoparticles. For membrane-bound and internalised aggregates, the wrapping and deformation of the membrane occurs either by tight packing of the nanoparticles or via the formation of U-structures, as can be seen in **Figs.7g**. Astonishingly, the morphological structures observed in these simulations replicate very closely the tubular structures observed in both conventional and STED confocal microscopy suggesting that the binding energy controls the membrane deformations and consequently how the cell processes vesicular carriers.

**Figure 7:**
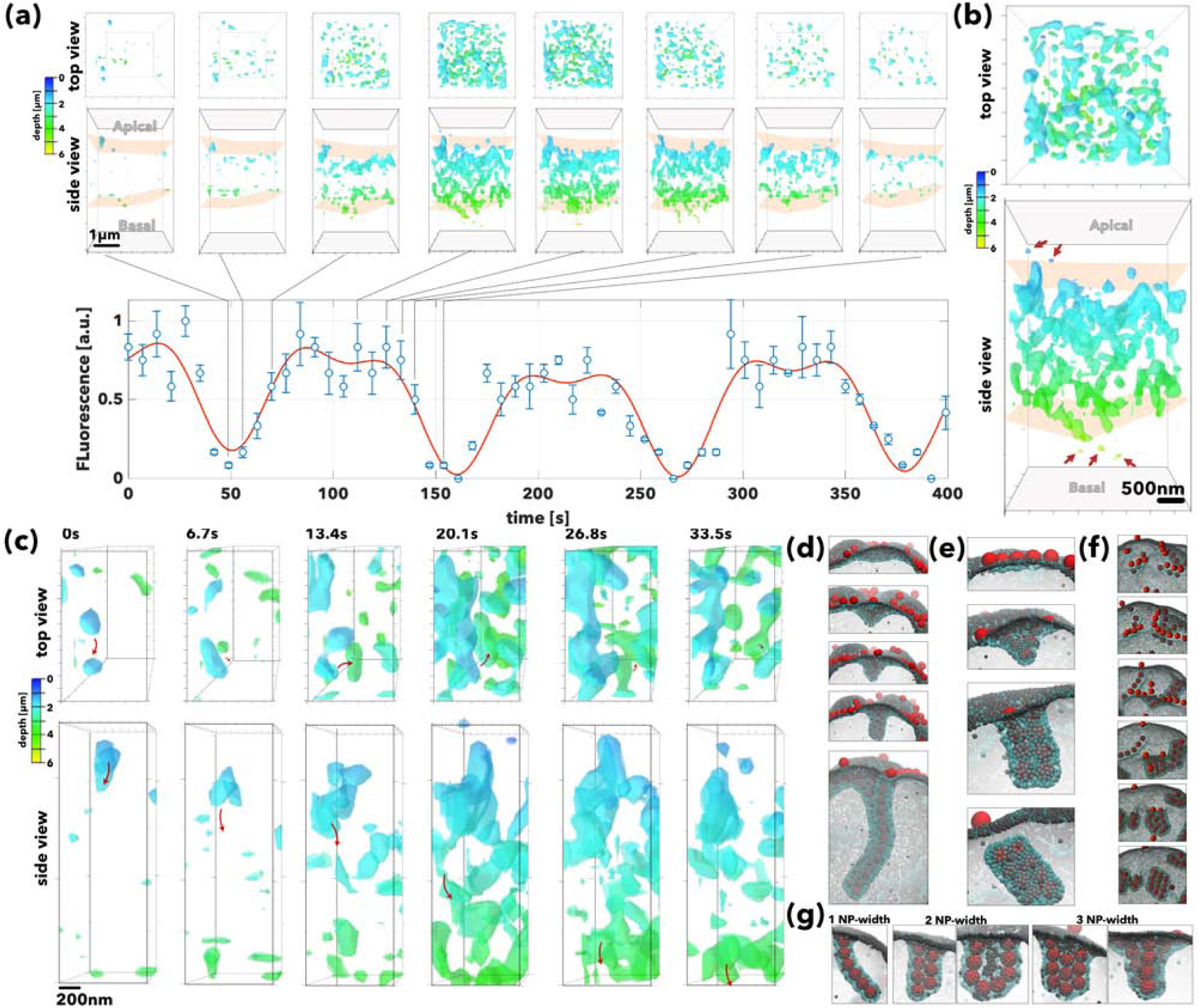
Super-resolution imaging of the tubular carriers.. 3D renderings showed as top and side views coloured coded as a function of the depth (*i.e*. the distance from apical to basal) of optical sections of brain endothelial cells (BECs) incubated with PtA2-loaded A_22_-P **(a)**. The 3D rendering was captured at a different time, and the normalised fluorescence measured across each section is plotted as a function of time. The 3D rendering at 120 seconds is shown enlarged as top and side view **(b)**, the arrows point at single A_22_-P particles while the structure emerged as a network of tubules. Close-up detail of top and side view of the same 3D rendering shows the evolution of the single tubule from apical to basal membrane showing the different stages of tubulation, fission from the apical membrane, transport and fusion to the basal membrane **(c)**. The same tubulations were observed in molecular dynamics simulations. Here the anisotropic growth of the membrane emerges from the collapse of a tubular aggregate of a particle on the surface. The membrane buckling can occur in different ways depending on the cluster size of assembled nanoparticles leading to the formation of tubules **(d)**, short tubes **(e)**, or multiple assemblies tubules **(f)**. The final tubule can be thus one, two or three nanoparticles (NP) thick **(g)**.

## Conclusions

Altogether, our findings show a very detailed picture of how LRP1 controls trafficking across BECs. We first demonstrate that this receptor is processed by both endo-lysosomal compartments as well as by tubular carriers that very likely distribute it between apical and basal membranes (**Fig.1**). We show that these tubular carriers are associated with the F-BAR domain syndapin-2 protein (**Fig. 2**), which probably functions as a structural stabiliser. By studying a model cargo targeting LRP1, we show that BECs sort the cargo depending on avidity (**Fig.3**) and this controls trafficking both at the binding/unbinding as well as membrane deformation. Not surprisingly, we observe that such a bias extends to intracellular trafficking. At high binding energy, the single cargo is internalised *via* a conventional endocytic pathway leading to lysosomal sorting and degradation, while the mid binding energy (“Goldilocks” avidity) leads to an unique pathway controlled by tubular carriers. These tubular carriers resemble morphologies previously reported, such as TEC [27, 26] and VVOs [28]. We thus show that avidity enables high efficiency of transport across BBB and becomes a discriminant for nutrients required for the brain or for endothelial cells themselves. Such a bias, in turn, alters the targeted receptor expression on BECs with the collective endocytosis and non-degradative tubular carrier pathway leading to LRP1 up-regulation while the single endocytic internalisation and endo-lysosomal pathway leads to down-regulation (**Fig.4**). The results, as demonstrated above, is that in the former the cargo is shuttled across efficiently while in the latter is degraded and possibly consumed by the same endothelial cell. Moreover, the ability to image the tubular carriers, by using the avidity optimised synthetic carrier A_22_-P, allowed us to reconstruct their morphology with an unprecedented resolution both *in vitro* and *in vivo* showing the critical role of syndapin-2 in stabilising them (**Fig.5**) and confirming previous TEM reconstructions [27]. Most importantly, we report here the dynamic of tubular formation and occurrence as well as their distribution over the cell surface (**Fig.6**). To this extent, the use of super-resolution imaging in combination with molecular dynamics simulation presents a critical role of the cargo in templating the tubular formation and dynamics (**Fig.7**). In conclusion, we shed light on BEC transcytosis using LRP1 as the main actor, but considering that “Goldilocks” avidity effect was also reported for transferrin receptor [49, 50, 51] and GLUT1 [52], our findings might suggest similar mechanisms. Nevertheless, ultimately, we report here a clear path to the brain, which once optimised it allows high delivery efficiency.

## Acknowledgements

G.B. thanks the ERC for the starting grant (MEViC 278793) and consolidator award (CheSSTaG 769798), the EPSRC/BTG Healthcare Partnership (EP/I001697/1), EPSRC Established Career Fellowship (EP/N026322/1), EPSRC/SomaNautix Healthcare Partnership EP/R024723/1, and Children with Cancer UK for the research project (16-227). X.T. and G.B. thank that Anhui 100 Talent program for facilitating data sharing and research visits. A.D.C and L.R. acknowledge the Royal Society for a Newton fellowship and the Marie Skłodowska-Curie Actions for a European Fellowship.

## Author contributions

X.T. prepared and characterised POs, performed all the fast imaging both in conventional and STED microscopy, setting up the initial BBB model, encapsulated the PtA2 in POs, and supervised the PtA2-PO animal work. D.M.L. prepared and characterised POs, performed all the permeability studies, PLA assays, WB and associated data analysis, part of the colocalisation assays and performed experiments with the shRNA for knockdown of syndapin-2. E.S. prepared and characterised POs, performed part of colocalisation assays, and Cy7-labelled PO animal experiments. S. N. prepared and characterised POs and performed part of the colocalisation and inhibition assays. G.F. designed, performed and analysed the agent-based simulations of transcytosis. J.F. designed the image-based algorithm to analyse the PLA data. D.L.M. prepared and characterised POs and helped with Cy7-labbelled PO animal experiments. A.A. performed TEM imaging of the POs. A.P. and A.D.C. synthesised the dye-, peptide-functionalised and pristine copolymers. M.V., L.K. and A.S. designed, performed and analysed the molecular dynamics simulations. Z.Z supervised and supported STED imaging. P.X., B.F. and Y.T. synthesised and characterised the PtA2 compound. L.L. performed some of the animal work. L.R. supported and helped with the BBB characterisation. G.B. analysed all fast imaging, supervised and coordinated the overall work. X.T., D.M.L., E.S and G.B. wrote the manuscript.

## Methods

### Materials

Polymers were obtained through ATRP synthesis as previously reported [36]. bEnd3 (CRL-2299), Dulbecco’s Modified Eagle’s Medium (DMEM) and FluoroBrite DMEM were obtained from ATCC. Foetal bovine serum (FBS), penicillin/streptomycin, phosphate-buffered saline (PBS, pH 7.4), 0.25% trypsin-EDTA, rat tail collagen I, 3-5 kDa fluorescein isothiocyanate-dextran (FITC-dextran) and 65-85 kDa tetramethylrhodamine isothiocyanate (TRITC-dextran) were obtained from Sigma-Aldrich. Transwell permeable polyester membranes (400 nm pore, 1.12 cm^−2^) were obtained from Corning Inc. EVOM2 Epithelial Voltohmmeter with STX3 electrodes was purchased from Word Precision Instruments. Paraformaldehyde (PFA), Triton X-100, normal horse serum, FITC-conjugated lectin, PLA probe anti-rabbit PLUS, PLA probe anti-mouse MINUS, Duolink detection reagent orange, radioimmunoprecipitation (RIPA) buffer, Tween-20, dextran (60-76 kDa), dynasore, N-ethylmaleimide (NEM), methyl-ß-cyclodextrin (CD), bovine serum albumin (BSA) and cholesterol quantification kit were also obtained from Sigma-Aldrich. Vectashield Mounting Media was obtained from Vector Labs. Protease Inhibitors, BCA protein assay kit and Laemmli sample buffer (x4) were purchased from Biorad. Angiopep-2 was obtained from GenScript. Puromycin dihydrochloride, gentamicin, CellMask*™* plasma membrane deep red, 4’,6-diamidino-2-phenylindole (DAPI) and Leica standard immersion oil were obtained from Thermo Fisher Scientific. All antibodies used are listed above. Polybrene, syndapin-2 shRNA and control shRNA lentiviral particles were purchased from Santa Cruz Biotechnology.

### Animal experiments

All animal studies were carried out according to the guidelines of ARRIVE (Animal Research: Reporting of *in vivo* Experiments) under licence from the UK Home Office (Scientific Procedures Act 1986) and approved by the University College London ethical review committee. Other set of animal experiments were carried out according to the national regulations and approved by the animal experiments ethical committee of School of Pharmaceutical Sciences and School of Chinese Medicine, Southwest University. In all experiments, animals were housed in a controlled temperature room with regular alternating cycles of light and darkness.

### Cell culture

Mouse brain endothelial cells bEnd3 were used between passage 20-30. bEnd3 were grown in DMEM supplemented with 10% (v/v) FBS and 100 IUL mL^−1^ penicillin/100 mg mL^−1^ streptomycin. Cells were maintained at 37 °C in an atmosphere of 5% CO2. For subculture, bEnd3 cells were washed twice with PBS, incubated with 0.25% trypsin-EDTA for 3 minutes, centrifuged and resuspended in fresh media. Media was changed every 2-3 days.

### *In vitro* BBB Model

To form a polarised confluent brain endothelial cell monolayer, bEnd3 were seeded at a density of 25,000 cells cm^−2^ in collagen-coated polyester membranes. bEnd3 were grown for 3 days in complete DMEM medium containing 10% (v/v) FBS, and then switched to serum-free media in the basal side of the transwell membrane for another 3 days. On day 6, transendothelial resistance was measured using an EVOM2 and the expression of PECAM, claudin-5 and ZO-1 was assessed by immuno-fluorescence. Dextrans (3-5 and 65-85 kDa) permeability across the endothelial monolayers was also assessed. A detailed description of the permeability assays can be found in Supporting Information.

### Polymersome preparation

A description of PO preparation and characterisation *via* dynamic light scattering (DLS) and transmission electron microscopy (TEM) can be found in the Supporting Information section.

### Immuno-fluorescence

Polarised bEnd3 monolayers either untreated or treated with A_22_-P (500 *μ*g mL^−1^) were washed twice with PBS, fixed in 4% (w/v) PFA for 15 minutes, permeabilised with 0.1% (w/v) Triton X-100 in PBS for 10 minutes and incubated with 5% (w/v) BSA in PBS for 1 hour at room temperature. Afterwards, cell monolayers were incubated with primary antibodies diluted in 1% (w/v) BSA and 0.01% (w/v) Triton X-100 in PBS overnight at 4 °C, followed by washing with PBS and incubation with the corresponding secondary antibodies for 2 hours at room temperature. Nuclei was counterstained by incubation with DAPI for 10 minutes. Transwell membranes were excised using a scalpel and mounted on co-verslips with Vectashield Mounting Media. Coronal brain sections were obtained from adult C57BL/6J (4 months old) mice. Briefly, brain sections were incubated in 20% (v/v) normal horse serum in PBS containing 0.3% (w/v) Triton X-100 for 2 hours at room temperature under gentle agitation followed by incubation with primary antibody antisyndapin-2 overnight at 4 °C. Sections were washed with PBS, incubated with the corresponding secondary antibody and FITC-conjugated lectin (1:200) for 2 hours and washed with PBS. Brain sections were mounted on glass slides in Vectashield Mounting Media. List of antibodies in Supporting Information.

### Western blot

Polarised bEnd3, either untreated or treated with angiopep-2 (1.75 nM), A_22_-P or A_110_-P (500 *μ*g mL^−1^) for 0.25, 1 and 2 hours, were washed twice with PBS and RIPA buffer containing protease inhibitors (1:50) was added directly to the membranes and left on ice for 1 hour. Cells were collected, centrifuged and the supernatant was collected for Western Blot analysis. Protein levels in the cell lysates were determined using BCA Protein Assay Kit. Lysates were mixed with Laemmli sample buffer and proteins (10 *μ*g) were separated on 10% SDS polyacrylamide gels and transferred to polyvinylidene difluoride (PVDF) membranes. Membranes were blocked with 5% (w/v) non-fat milk in Tris-buffered saline (TBS) containing 0.1% (w/v) Tween-20 (TBS-T) for 1 hour and then incubated with a rabbit monoclonal antibody to LRP1 overnight at 4 °C. After washing with TBS-T, the membranes were incubated with a secondary antibody for 2 hours at room temperature and imaged using an Odyssey CLx (LI-COR Biosciences). The membranes were further probed for GAPDH as a loading control.

### Proximity ligation assay

Polarised bEnd3 (untreated or treated with angiopep-2 (1.75 nM), A_22_-P or A_110_-P (500 *μ*g mL^−1^) for 0.25, 1 and 2 hours) were washed twice with PBS, fixed in 4% (w/v) PFA in PBS for 15 minutes and permeabilised with 0.1% (w/v) Triton X-100 in PBS for 10 minutes. For the PLA assay, the Duolink probes and detection reagents were used according to the supplier’s instructions. Briefly, monolayers were incubated with Duolink blocking solution for 1 hour at 37 °C, and then incubated with two antibodies targeting the proteins of interest (one being LRP1 and the other one of the proteins relevant for transcytosis) overnight at 4 °C. Following incubation with primary antibodies, cells were incubated with the Duolink PLA probes (anti-rabbit and anti-mouse) for 1 hour at 37 °C, washed and incubated with Duolink ligase and polymerase for 30 and 100 minutes, respectively. Nuclei were stained by adding DAPI for 10 minutes. Membranes were mounted in glass coverslips using Vectashield Mounting Media. List of antibodies in Supporting Information.

PLA data were quantitatively analysed using a Python script based on Trackpy modified for identification of particles with high polydispersity in the direction of objective translation (‘z’). A median filter and a low-pass Gaussian filter were applied to each slide in the confocal stack to remove and large scale features due to residual channel cross-talk. Local maxima were then identified, and maxima in the z-direction corresponding to the same particle were grouped by hierarchical clustering using the Nearest Point Algorithm as implemented in scipy [57]. For quantitative fluorescence measurements, a correction was applied to account for the variation in photon dose per unit volume as a function of voxel size by multiplying signal intensity by a factor of 1/(δ_*x*_ δ_*y*_ δ_*z*_), where δ_*x*_, δ_*y*_, and δ_*z*_ were the voxel dimensions in the x, y, and z direction, respectively.

### Permeability Assays

To assess permeability across polarised bEnd3, Cy7-labelled A_*L*_-P with *L* = 0, 22, 36, 56, 110 and 220 (100 *μ*g mL^−1^) and FITC-angiopep-2 (10 *μ*g mL^−1^) were added to the apical side of the Transwell membrane and incubated at 37 °C. Samples were collected from the basal side and fresh medium was added to replace the volume. Fluorescence intensity of the Cy7-labelled A_*L*_P or FITC-angiopep-2 was measured in black 96-well plates using a Spark Multimode microplate reader (Tecan). Apparent permeability was calculated using **Equation 1**.

### *In vivo* biodistribution of Cy7-labelled POs

Healthy adult C57BL/6J female mice were injected *via* tail vein with 100*μ*L of either Cy5- and Cy7-labelled A_*l*_-P with *L* = 0, 22 or 110 at a concentration of 4 mg mL^−1^ or with free Cy5-angiopep-2. Within two hours from the administration, mice were anaesthetised with isoflurane and imaged using an IVIS (PerkinElmer). Animals were then perfused with 50 mL of PBS, brains were collected, imaged again using IVIS and then snap frozen in liquid nitrogen. To quantify brain accumulation, the cerebellum was removed. Cerebrum was weighed, PBS was added and manually homogenised with the addition of 3 volumes of 30% (w/v) dextran (64-74 kDa). Then, samples were centrifuged 7400 *g* for 20 minutes, which results in the separation into fractions: capillary-depleted fraction (*i.e*., parenchyma), dextran, and capillary-enriched fraction (*i.e*., vessels). Parenchyma was resuspended and added to a black 96-well plate. Fluorescence of the POs was measured in a Spark Multimode microplate reader. Sample fluorescence readings were normalized to the ones obtained from the mice injected with PBS. Positive control was POs spiked at a known concentration in the homogenates (*n* = 3). Normalised fluorescence readings were converted to POs mass and that was then converted into the percentage of ID per gram (% ID/g) of tissue.

### *In vivo* biodistribution of Pt2A-loaded POs and histological analysis

A detailed description of the synthesis and characterisation of PtA2 and preparation of PtA2-loaded POs can be found in Supporting Information. For the *in vivo* biodistribution of PtA2-loaded POs, four weeks old Kunming mouse were injected *via* tail vein with POs (200 *μ*L at 1 mg mL^−1^). At given time points, mice were culled with an overdose of isoflurane, perfused with PBS and the organs fixed in a solution of 4% (w/v) PFA in PBS. The organs were further fixed with 4% (w/v) PFA at 4 °C for 24 hours and dehydrated in a solution of 30% (w/v) sucrose in PBS for more than 48 hours. Next, the organs were weighed and digested in 60% (v/v) nitric acid at room temperature for 24 hours. Each organ was diluted in ultra-pure water to achieve a final volume of 10 mL containing 3% (v/v) nitric acid. The concentration of platinum in each organ was determined using an inductively coupled plasma mass spectrometer (ICP-MS, Thermo Fisher). Alternatively, the perfused brains were extracted, dura mater removed and post-fixed for 7 hours in 4% PFA at 4 °C. Then, the fixed brains were immersed in 20% (w/v) sucrose in PBS overnight at 4 °C for cryoprotection. Fixed brains were cut using a cryostat (Leica 1950) at 20*μ*m in the coronal plane and mounted onto glass slides. Sections were initially washed three times in PBS for 5 minutes at room temperature and pre-incubated for 1 hour in a blocking buffer consisting of 2% (v/v) goat serum, 1% (w/v) BSA in 0.1% (v/v) Triton X-100 in PBS (PBS-T). After washing with PBS, processed sections were incubated with primary antibodies diluted in PBS for 1 hour at room temperature. Sections were then washed four times in PBS for 5 minutes each and incubated for 2 hours at room temperature using the appropriate secondary antibodies. For blood vessel labelling, mice were initially injected intravenously with FITC-labelled lectins (200 *μ*L of 500 *μ*g mL^−1^f) at 5 minutes before culling.

### Confocal microscopy and imaging

For live kinetics, bEnd3 monolayers were incubated with CellMask*™* Deep Red plasma membrane staining for 30 minutes, rinsed with PBS three times and immersed in a FluoroBrite DMEM medium supplemented with 10% (v/v) FBS and gentamicin (5 mg mL^−1^). Subsequently, Cy5-labelled A_22_-P were added (1 mg mL^−1^) into the apical side of the transwell and incubated for 1 to 2 hours at 37 °C in 95% air and 5% CO2. Images were acquired using a Leica TCS SP8 confocal microscope equipped with Diode 405, Argon, DPSS 561 and HeNe633 lasers. Images were acquired via sequential scan to reduce fluorophore bleed-through. For live-cell imaging, an incubator at 37 °C and 5% CO2 connected to the unit was used, and allowed to stabilise for 1 hour before imaging. For live-cell 3D scanning, images were acquired with a 10X 0.3NA objective in resonant scanning mode with a speed of 700Hz and a resolution of 128×512 pixels. For fixed cell imaging, images were acquired with a 63x oil immersion objective at 400Hz and 512×512 pixels. Leica SP8 AFS microscope software was used to generate 3D projections from z-stacks. The same software was used for analysis of POs fluorescence in transwell z-stacks and normalisation of fluorescence intensity. ImageJ was used for analysis of fluorescence intensity in the xy plane. Colocalisation analysis to obtain Pearson’s correlation coefficient *r* was done using the plug-in for colocalisation on ImageJ.

### Stimulated emission depletion microscopy imaging

Stimulated emission depletion (STED) and Fast-STED super resolution imaging experiments were done using a Leica DMi8 confocal microscopy equipped with a Leica TCS SP8 STED-ONE unit. PtA2-loaded POs were excited with a 405 nm laser and emission was collected at 550 - 580 nm with donut laser at 595 nm. Images were collected using HyD reflected light detectors (RLDs) with 2048×2048 pixel and x100 scanning speed. For Fast-STED imaging, a RESONANT model was applied to minimise the 3D scanning time. Transwell membranes were imaged with a 100x lens. 3D Images were recorded after POs were added into the apical side (thickness of 8 *μ*m, interval = 0.2 *μ*m, time interval 5.0 – 8.0 seconds). STED and Fast-STED micrographs were further processed using ‘deconvolution wizard’ function from Huygens Professional software under authorised license. The area radiuses were estimated under 0.03 *μ*m with exclusion of 200 absolute background values. Maximum iterations was 10-times and signal-to-noise ration of 30 was applied, with quality threshold at 0.005. Other settings were the “optimised” iteration mode and the “automatic” brick layout.

### Statistics

The results are expressed as mean ± standard deviation (SD). Comparisons between groups were obtained by One-Way ANOVA using Dunnett’s post hoc test in the comparison to a control or Tukey’s for multiple comparisons between groups in GraphPad Prism 7.03. Significance level of *P* < 0.05 was considered statistically significant.

## 1 Supporting Information

### Characterisation of the *in vitro* BBB Model

bEnd3 were seeded at a density of 25,000 cells cm^−2^ in collagen-coated polyester membranes. Cells were grown for 3 days in complete DMEM medium containing 10% (v/v) FBS, and then switched to serum-free media in the basal side of the transwell membrane for another 3 days. On day 6, expression of PECAM, claudin-5 and ZO-1 was assessed by immuno-fluorescence.Dextrans (3-5 and 65-85 kDa) permeability across the endothelial monolayers was also assessed. Briefly, FITC- and TRITC-dextrans (1 mg mL^−1^) were added to the apical side of the transwell membrane containing media in the basal side. At specific time-points, a sample was collected from the basolateral side and fresh media was used to replace the volume. Fluorescence of FITC and TRITC was measured in black 96 wellplates using Spark Multimode Microplate Reader (Tecan). Apparent permeability was calculated according to Equation 1 (**Fig.S1**).

### Preparation and characterisation of POs

Poly(ethyleneglycol) methyl ether methacrylate (POEGMA)_20_-poly(2-(diisopropylamino)ethyl methacrylate) (PDPA_100_) copolymer was prepared according to a previously published procedure [36]. Angiopep-2- and Cy5-/Cy7-labelled POEGMA_20_-PDPA_100_ were prepared according to established protocols [36]. Angiopep-2 (0-12%mol), Cy5- or Cy7-(10%mol)-functionalised POs were prepared either by pH or solvent-switch approach. Briefly, in a pH-switch, copolymer (20 mg) was dissolved in PBS at pH 2, followed by gradually increasing the pH to 6.0 using 0.5 M NaOH under magnetic stirring. At pH 6.0, angiopep-2-POEGMA_20_-PDPA_100_ was added to the copolymer solution and the pH was then increased to 7.4. In solvent-switch, copolymer (20 mg) was dissolved in tetrahydrofuran (THF), and to this polymer solution, PBS (pH 7.4) was added using a syringe pump at a flow rate of 2 *μ*L min^−1^ under stirring at 40 °C. A final volume of PBS (2.3 mL) was added. Once the volume of PBS was added, POs were dialysed against PBS (pH 7.4). In both approaches, POs were purified *via* a gel permeation chromatography on a size-exclusion column containing Sepharose 4B. Dynamic scattering (DLS) was used to assess average size and polydispersity of the POs by using a Malvern Zetasizer Nano ZS. POs were imaged by transmission electron microscopy (TEM) to assess size, surface topology and morphology using a JEOL microscope with a 100 kV voltage tension (**Fig.S2**).

### Synthesis and characterisation of Pt2A

Initially, FA2 compound was synthesised. Briefly, ammonium acetate (7.71 g, 0.10 mol), 4-methylbenzaldehyde (1.52 g, 0.10 mol), 1-tetralone (2.89 g, 0.02 mol) and glacial acetic acid (40 mL) were added to a 100 mL round-bottom flask and stirred for 24 hours at room temperature to obtain a burgundy solution. At the end of the reaction (monitored by thin layer chromatography), the resulting product was poured into an excess of water. The obtained solid was filtered by suction an purified by column chromatography using petroleum ether and ethyl acetate (10:1) as eluent. A white solid (5.03 g) was then obtained and further purified by recrystallising in ethyl acetate (yield of 89%). ^1^H-nuclear magnetic resonance (^1^H-NMR) was performed (400 MHz, CD3CN) showing the following peaks *δ* (ppm): 8.51 (*d,J* =7.7 Hz, 1H), 7.42 (*t,J*)=7.7 Hz, 1H), 7.35 (*t,J*)=7.4 Hz, 1H), 7.28 (*d,J*)=7.4 Hz, 4H), 7.07 (*d,J*)=8.4 Hz, 6H), 6.84 (*d,J*)=8.2 Hz, 2H), 3.45 (*q,J*)=7.0 Hz, 2H), 2.91-2.82 (m, 4H), 1.21 (*t,J*)=7.0 Hz, 4H). High resolution mass spectrometry (HRMS) electrospray ionisation mass spectrometry (ESI-MS) was also carried out, in which theroretical m/z is 373.49 and the one obtained was 374.19 ([M+H] +).

Once the FA2 compound was obtained, PtA2 was then synthesised. Briefly, FA2 (0.53 g, 1.3 mmol) and potassium tetrachloroplatinate(II) (0.50 g, 1.2 mmol) were mixed in acetic acid (150 mL). The liquid suspension was heated to 400 K for 72 hours under nitrogen atmosphere. A yellow-green residue was obtained after filtration. The intermediate, without further purification, was dissolved in dimethyl sulfoxide (5 mL) and refluxed for 30 minutes. Then, the reaction solution was treated with deionised water (50 mL). A yellow powder was obtained after filtration and dried under vaccum conditions. The purified product was obtained by column chromatography (neutral alumina, hexane:ethyl acetate)c = 5:1) as a yellow powder. Yield was found to be 53%. Final product was characterised by ^1^H-NMR (400 MHz, d6-DMSO) showing the distinctive peaks *δ* (ppm): 7.46 (*t,J* =8.1 Hz, 2H), 7.35 (*d,J* =7.8 Hz, 2H), 7.23 (*d,J* =8.0 Hz, 2H), 7.07 (*dd,J* =15.3, 8.0 Hz, 2H), 6.83 (*dd,J* =31.1, 7.5 Hz, 2H), 2.78 (*t,J* =7.6 Hz, 4H), 2.53 (*d,J* =5.0 Hz, 10H), 2.40 (s, 3H). Fourier-transform infrared spectroscopy (FTIR))was also performed (KBr, *v*, cm^−1^) exhibiting the peaks: 3433, 3037, 2912, 1603, 1567, 1390, 1310, 1244, 1112, 1016, 765, 685, 450. HRMS (ESI-MS) was performed with a calculated m/z of 644.69 and the m/z found of 645.15 ([M+H] +) (**Fig.S3**).

### Preparation of PtA2-loaded POs

PtA2 (1 mg) and copolymer (10 mg) were individually dissolved into a mixture of methanol and dichloromethane (5 mL) in separate vials. Both solutions were then mixed uniformly into a 10 mL flat-bottom glass bottle, and rotaevaporated at 40 ^°^C to obtain a thin film. The resulting film 24 was hydrated with PBS (5 mL) and left to stir at room temperature for at least 2 weeks. Morphology and PtA2 encapsulation were evaluated by TEM and fluorescence spectrophotometry, respectively.

### Transcytosis model

Nanoparticles were randomly seeded within the aqueous phase in the starting states. Nanoparticles in the aqueous phase moved according to Brownian motion with a time step of 0.00005 seconds. Dynamic viscosity was given as 0.00078 Pa ^−1^, which corresponds to the viscosity of the cell medium DMEM at a standard temperature of 37 ° (310 K) [1]. The boundaries of the transwell, the top of the aqueous phase and the edge of the cell layer were treated as reflective boundaries. The characteristic length for LRP1-angiopep-2 was estimated to be 6.5 nm based on the molecular weight of LRP1*β* (the extracellular chains that forms part of the LRP1 heterodimer) using the methods of Erickson [2, 3]. Nanoparticles in close enough proximity to the top of the cell layer are able to form bonds with the cell. Temporary ligand agents were created randomly outside of these nanoparticles to the required density as this method would be computationally less expensive than adding rotational diffusion and updating the positions of the ligands each iteration. Moreover, for the time step used for receptor binding (0.01 seconds) and the radius of the nanoparticles, the contact surface is effectively randomised each iteration by rotational diffusion so that recreation of the ligands would not adversely effect the robustness of the model compared to incorporating full rotational diffusion. Therefore, this method was adapted to give a simulation of binding and unbinding across discrete steps. Decuzzi and Ferrari investigated the binding of nanoparticles in a static linear flow, they gave the probability of a nanoparticle adhering as:

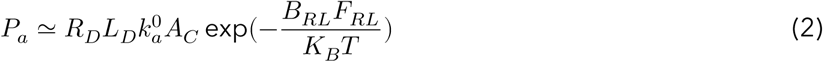

*R*_*D*_ is the receptor density on the cell surface, 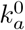 is the association constant at zero load per receptor-ligand pair and *A*_*c*_ is the contact area between nanoparticle and cell surface.

The interfacial contact surface area, the receptor density and the ligand density dictate the number of bonds between the particle and a cell. The target cell determines the receptor density whilst ligand density is an adaptable property of the nanoparticles. The interfacial surface area is the surface area of the nanoparticle that is within a set binding distance from the cell. The characteristic length of the receptor-ligand bond. We then build an agent-based model of nanoparticle-cell binding unit based on the transwell *in vitro* model of the BBB to investigate how adapting the ligand density and nanoparticle size can alter transcytosis efficiency (**Fig.S4**).

### Molecular dynamics simulations

A meshless solvent-free coarse-grained (CG) membrane model was coupled with molecular dynamics (MD) simulations to capture membrane topological changes and nanoparticle aggregation dynamics at the required length and time scales. In the model, the membrane was discretised into beads that each represent a CG membrane surface patch (**Fig.S5**). Different beads were used to represent a membrane surface patch with no receptors (‘inert’ bead) and a membrane surface patch with receptors (‘receptor’ bead). The beads self-assembled into a membrane and replicated biologically relevant properties using a soft-core pairwise inter-particle potential [4]. An equilibrated spherical membrane of 20162 membrane beads of diameter 1*σ* (where*σ*is the MD unit of length) was used in the simulations. Typically, nanoparticles were represented as beads of diameter 4*σ* in the model. 105 nanoparticles were randomly distributed in a spherical shell surrounding the membrane with a radius *R*_*M*_ + 6*σ*, where *R*_*M*_ is the undeformed average radius of the membrane. A minimum distance of 5*σ* between the centers of the nanoparticles was enforced in this initial distribution. The repulsive branch of a 12-6 Lennard-Jones potential was used for the volume exclusion of the nanoparticles. The nanoparticle-receptor affinity was modelled using the attractive branch of a 12-6 Lennard-Jones potential. This attractive potential between the nanoparticles and ‘receptor’ membrane beads was cutoff at *r*_*c*_ = 3.75*σ*. Therefore for each *r* < *r*_*c*_, the 12-6 Lennard-Jones potential between nanoparticle and receptors is:

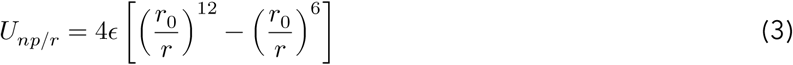

where *ϵ* is the depth of the potential well, *r*_0_ the distance at which the potential is zero, and *r* the distance between particles.

For the membrane mechanics, we adopted a meshless model. Meshless models can capture membrane topological changes and time dynamics naturally when coupled with MD simulations. However, a careful choice of potential between membrane beads is essential to ensuring a faithful replication of biologically relevant membrane properties. We employed the soft-core pairwise inter-particle potential developed by Yuan, et. al. [4]. A 4-2 Lennard-Jones (LJ) type potential was used for the repulsive branch of the potential that ensured particle volume exclusion and a cosine function potential was used for the attractive branch of the potential for driving membrane self-assembly. The potential was used for interactions between and among membrane and receptor beads. The molecular dynamics simulations were carried out in the NVE ensemble, where *N* is the total number of system particles, *V* is the volume of the simulation box and *E* is the total energy of the system. A Langevin thermostat was applied to the system components to model interactions with an implicit background solvent. The simulations were typically carried out for 2,000,000 time steps, with a time step of 0.01ŁŁ (where ŁŁ is the MD unit of time). The simulations were implemented with the LAMMPS package [5].

### Immuno-fluorescence

Polarised bEnd3 were washed twice with PBS, fixed in 4% (w/v) paraformaldehyde (PFA) for 15 minutes, permeabilised with 0.1% (w/v) Triton X-100 in PBS for 10 minutes and incubated with 5% (w/v) BSA in PBS for 1 hour at room temperature. Afterwards, cell monolayers were incubated with primary antibodies diluted in 1% (w/v) BSA and 0.01% (w/v) Triton X-100 in PBS overnight at 4 °C, followed by washing with PBS and incubation with the corresponding secondary antibodies for 2 hours at room temperature. Nuclei was counterstained by incubation with DAPI for 10 minutes. Transwell membranes were excised using a scalpel and mounted on coverslips with Vectashield Mounting Media. Antibodies used in our studies are listed on **Table 1**.

**Table S1.** List of antibodies.

| Antibody | Dilution | Supplier, Catalogue Number |
| --- | --- | --- |
| Goat polyclonal to PECAM-1 | 1:200 | RD Systems, AF3628 |
| Rabbit polyclonal to claudin-5 | 1:100 | Abcam, ab15106 |
| Rabbit polyclonal to ZO-1 | 1:100 | Abcam, ab221547 |
| Mouse monoclonal to LRP1 | 1:1000 | Abcam, ab28320 |
| Rabbit monoclonal to LRP1 † | 1:1000 | Abcam, ab92544 |
| Mouse monoclonal to GAPDH † | 1:1000 | Abcam, ab8245 |
| Rabbit polyclonal to syndapin-2 † | 1:400 | Abcam, ab37615 |
| Rabbit polyclonal to caveolin-1 | 1:100 | Sigma-Aldrich, C4490 |
| Rabbit polyclonal to clathrin | 1:1000 | Abcam, ab21679 |
| Rabbit polyclonal to dynamin-2 | 1:300 | Abcam, ab3457 |
| Rabbit polyclonal to EEA-1 | 1:100 | Abcam, ab2900 |
| Rabbit polyclonal to Rab5 | 1:100 | Abcam, ab13253 |
| Rabbit monoclonal to Rab7 | 1:100 | Abcam, ab137029 |
| Rabbit polyclonal to Rab11 | 1:100 | Abcam, ab3612 |
| Rabbit polyclonal to LAMP-1 | 1:100 | Abcam, ab24170 |
| Rabbit polyclonal to β-actin | 1:100 | Abcam, ab8227 |
| Rabbit monoclonal to myosin | 1:200 | Abcam, ab92721 |
| Rabbit polyclonal to α-tubulin | 1:100 | Abcam, ab18251 |
| Rabbit polyclonal to dynein | 1:200 | Abcam, ab236594 |
| Rabbit polyclonal to kinesin | 1:500 | Abcam, ab62104 |
| Alexa Fluor 488 goat anti-mouse IgG | 1:500 | Biolegend, 405319 |
| Alexa Fluor 647 donkey anti-rabbit IgG | 1:500 | Biolegend, 406414 |
| Alexa Fluor 488 donkey anti-rabbit IgG | 1:500 | Biolegend, 406416 |
| Alexa Fluor 488 donkey anti-goat Ig | 1:500 | Abcam, ab150129 |
| Dylight 800 goat anti-mouse IgG † | 1:5000 | Thermo Fisher Scientific, SA535521 |
| Dylight 800 goat anti-rabbit IgG † | 1:5000 | Thermo Fisher Scientific, SA535571 |

† Antibodies used for western blot.

### Competition assay

For the compeptition assay, FITC-angiopep-2 was dissolved in ultrapure water at 1.75 pM, which is the concentration equivalent to the ligand functionalisation inA_22_-P. Angiopep-2 or Cy5-labelled A_22_-P were added separately or together to the apical side of the transwell and cells were incubated at 37 °C. After 10 and 60 minutes, cells were washed twice with PBS and fixed in 4% PFA. Transwell membranes were excised with a scapel and mounted on coverslips using VectaShield Mounting Medium with DAPI for confocal imaging. Quantification of intracellular fluorescence of Cy5-labelled A_22_-P or FITC was performed *via* ImageJ normalising fluorescence to the number of cell nuclei (DAPI) (**Fig.S6**).

### Small molecule pharmacological inhibitors

Polarised bEnd3 cells were incubated with CellMask*™* Deep Red for 10 minutes at 37 °C and then rinsed with PBS. Dynasore (40 *μ*M) was added to the cell media and pre-incubated for 10 minutes at 37 °C, followed by the addition of A_22_-P (100 *μ*g mL^−1^) for 60 minutes. After incubation, cells were washed and media was replaced with serum free FluoroBrite DMEM. For the dynasore recovery experiments, cells were washed with PBS at pH5 before the addition of the imaging media. In the inhibition experiments with N-ethylmaleimide (NEM), cells were incubated with NEM (0.5 mM) for 5 minutes before the addition of the A_22_-P (100 *μ*g mL^−1^) for 60 minutes. Live-cell imaging was performed using Leica TCS SP8 confocal microscope (**Fig.S7**).

### Membrane cholesterol depletion

Prior to the cholesterol depletion assay, media of polarised bEnd3 was changed to serum-free DMEM. Methyl-*β*-cyclodextrin (CD, 10 mM) was added either to the apical or basal side of the transwell and cells were incubated for 15 minutes at 37 °C. Apical and basal media was collected and used for the quantification of free cholesterol. A_22_-P (100 *μ*g mL^−1^) were added to the apical side of the transwell and incubated for 60 minutes at 37 °C. Afterwards, cells were washed twice with PBS and fixed in 4% PFA for 15 minutes, followed by incubation with 0.3% (w/v) Triton X-100 and 10% BSA in PBS for 30 minutes. Cells were incubated with anti-caveolin-1 antibody for 2 hours at room temperature followed by incubation with an appropriate secundary antibody. Cholesterol quantification in the media was performed by using a colorimetric cholesterol quantification kit according to the supplier’s instructions (**Fig.S8**).

### Lentiviral shRNA silencing

Short hairpin RNA (shRNA) lentiviral particles for syndapin-2 were used according to supplier’s instructions. Briefly, bEnd3 cells were seeded on a 6-well plate at a density of 100,000 cells per well and grown overnight. At 50% of confluence, cells were treated with the shRNA lentiviral particles in DMEM supplemented with polybrene (5 *μ*g mL^−1^) and then, incubated overnight. On the next day, the media was replaced, and cells were further incubated for 2 days. Stable clones expressing shRNA were selected by puromycin (5 *μ*g mL−1) and silencing of syndapin-2 was then confirmed by Western blot. Control shRNA lentiviral particles were used as a negative control (**Fig.S10**).

### Transmission electron microscopy of brain tissue

Brains from mice treated with PtA2-loaded POs were cut into 1×1×1 mm size cubes. Brain tissue was fixed in fresh 3% (v/v) glutaradehyde in phosphate buffer (0.1 M) overnight at 4 °C, and then washed twice in phosphate buffer (0.1 M) at 4 °C at 30 minutes intervals. Samples were dehydrated through a series of ethanol incubations: 75% (15 minutes), 95% (15 minutes), 100% (15 minutes) and 100% (15 minutes) and then placed in an intermediate solvent (propylene oxide) for 2 changes of 15 minutes. Resin infiltration was obtained by placing the specimens in a 50:50 mixture of propylene/araldite resin leaving the samples with the mixture overnight at room temperature. Afterwards, speciments were moved into Araldite resin for 6-8 hours at room temperature with a change of resin after 3-4 hours. Finally, specimens were embedded in fresh Araldite resin for 48-72 hours at 60 °C. Semi-thin sections of approximately 0.5 *μ*m were cut on a Leica Ultramicrotome and stained with 1% Toluidine blue in Borax. Ultra-thin sections of 70-90 nm tick were cut on a Leica Ultramicrotome and stained for 25 minutes with saturated aqueous uranyl acetate followed by staining with Reynold’s lead citrate for 5 minutes. Sections were examined using FEI Tecnai TEM at an accelerating voltage of 80kVv. Electron micrographs were taken using a Gatan digital camera (**Fig.S11**).

[1] Rommel G Bacabac, Theo H Smit, Stephen C.Cowinc, Jack J W A Van Loon, Frans T M Nieuwstad, Rob Heethaar, and JennekeKlein-Nulenda. Dynamic shear stress in parallel-plate flow chambers. *Journal of Biomechanics*, 38: 159-167, January 2005. [2] Renad N Alyaudtin, Andreas Reichel, Raimar Lobenberg, Peter Ramge, Jorg Kreuter, and David J Begley. Interaction of poly(butylcyanoacrylate) nanoparticles with the blood-brain barrier in vivo and in vitro. *Journal of Drug Targeting*, 9(3):209-211. January 2001. [3] Ulrike Schroder and Bernhard A Sabel. Nanoparticles, a drug carrier system to pass the blood-brain barrier, permit central analgesic effects of i.v. dalargin injections. *Brain Research*, 710(1-2): 121-124. February 1996. [4] Hongyan Yuan, Changjin Huang, Ju Li, George Lykotrafitis, and Sulin Zhang. One-particle-thick, solvent-free, coarse-grained model for biological and biomimetic fluid membranes. *Physical Review E*, 82: 011905. July 2010. [5] Steve Plimpton. Fast Parallel Algorithms for Short-Range Molecular Dynamics. *Journal of Computational Physics*, 117(1): 1-19. March 1995.

---

**Figure S1:**
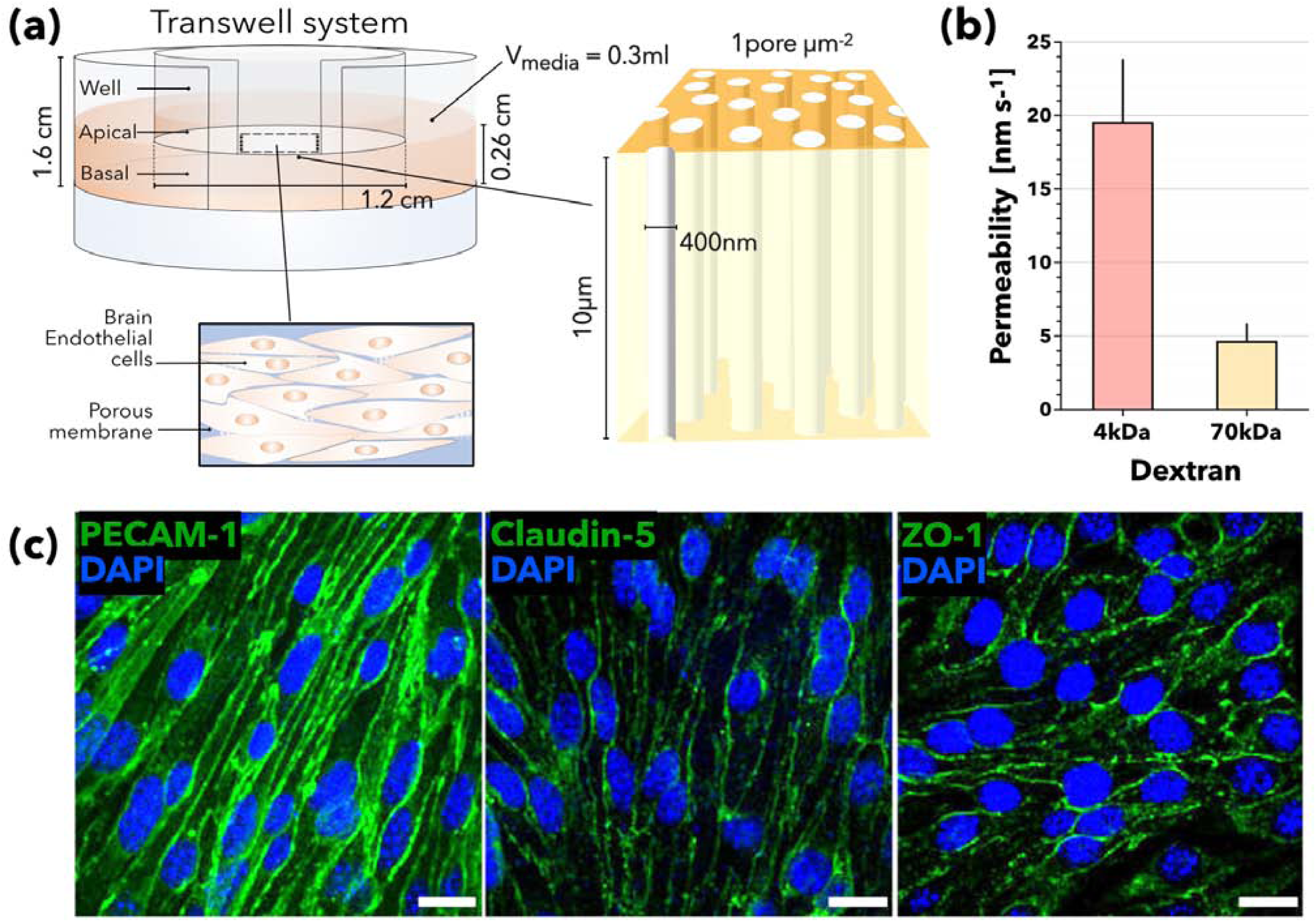
Characterisation of the *in vitro* BBB model. Schematic of the *in vitro* model of BBB used to assess transcytosis **(a)**. Bar chart showing the apparent permeability coefficient (*P*) of 4 and 70 kDa dextrans across bEnd3. Data represented as mean ± SD (*n* = 9) **(b)**. Confocal images of polarised bEnd3 showing expression of PECAM-1, claudin-5 and ZO-1. PECAM-1, claudin-5 and ZO-1 are shown in green and cell nuclei stained with DAPI (blue). Scale: 20 nm **(c)**.

**Figure S2:**
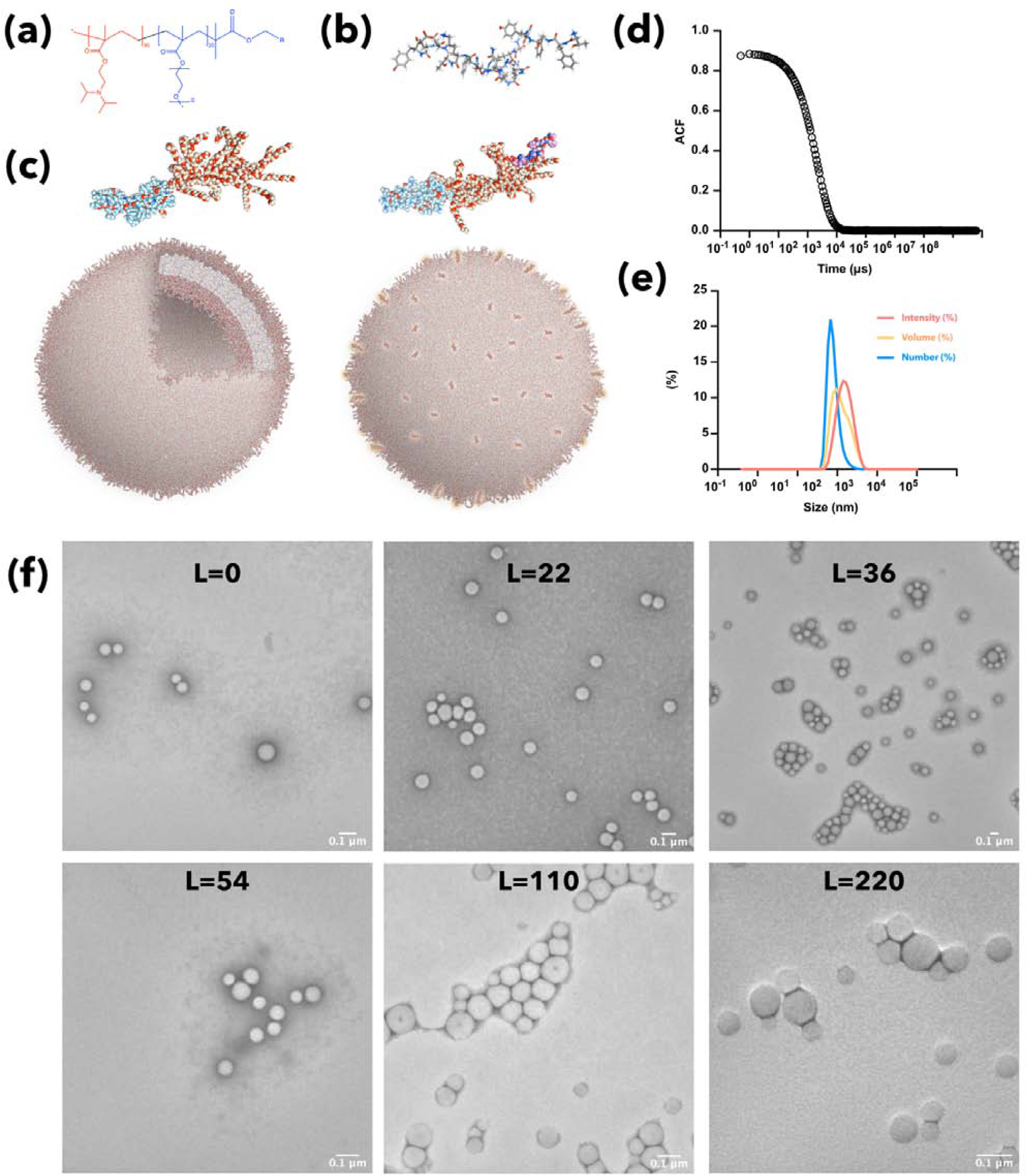
Characterisation of A_*L*_-POs. Schematic representation of POEGMA_20_-PDPA_100_ **(a)**, angiopep-2 **(b)**, and self-assembled pristine (left) and A_*L*_-POs (right) **(c)**. Representative correlogram **(d)** and histogram showing size distribution expressed as percentage of intensity (red), volume (yellow) and number (blue) **(d)** of the A_*L*_-POs. Data obtained by dynamic light scattering. Transmission electron micrographs of A_*L*_-P with *L* = 0, 22, 36, 54, 110 and 220.

**Figure S3:**
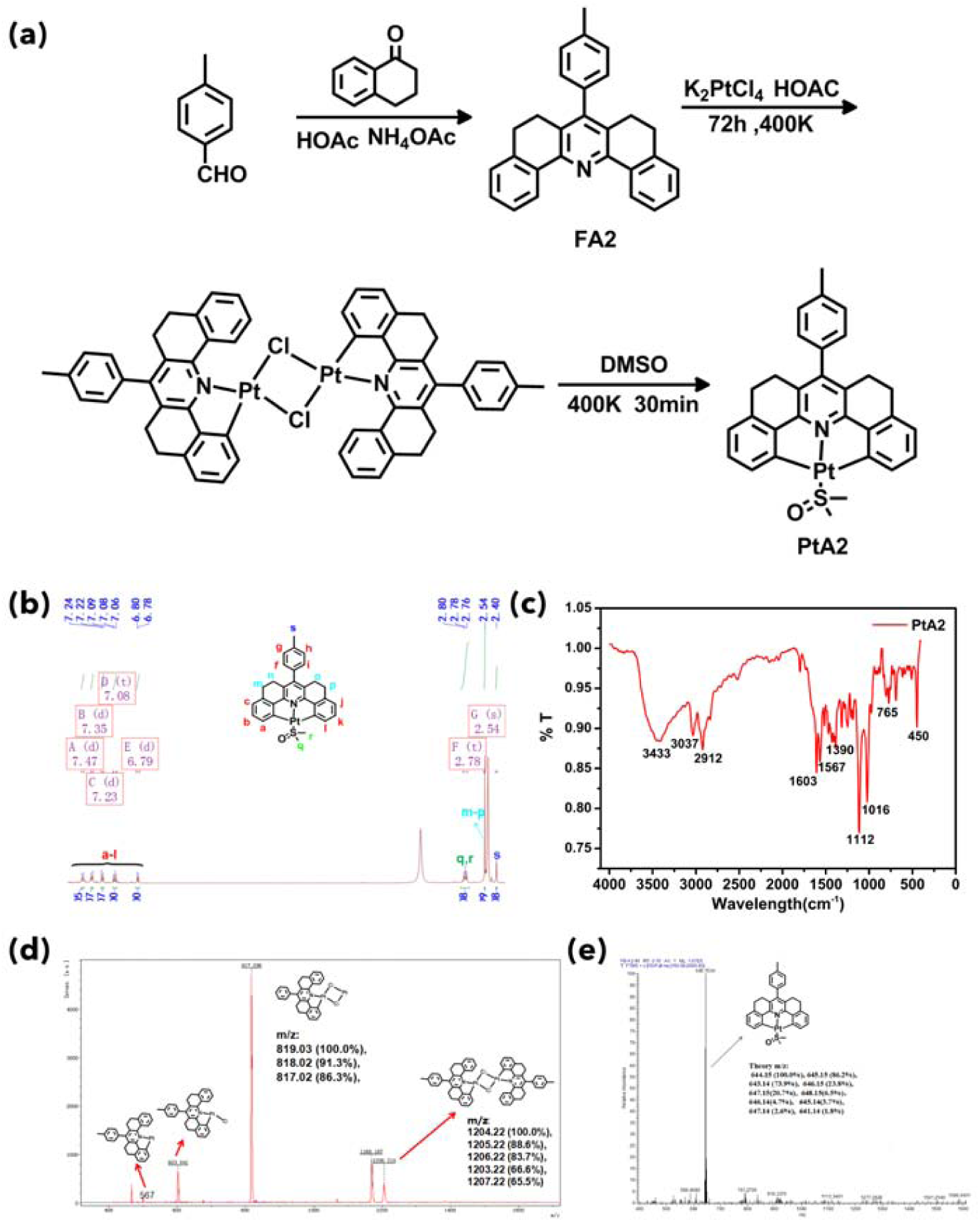
Characterisation of PtA2. A schematic representation of the synthesis of PtA2 **(a)**. ^1^H-NMR **(b)**, FTIR **(c)** and ESI-MS **(d,e)** spectra of purified PtA2.

**Figure S4:**
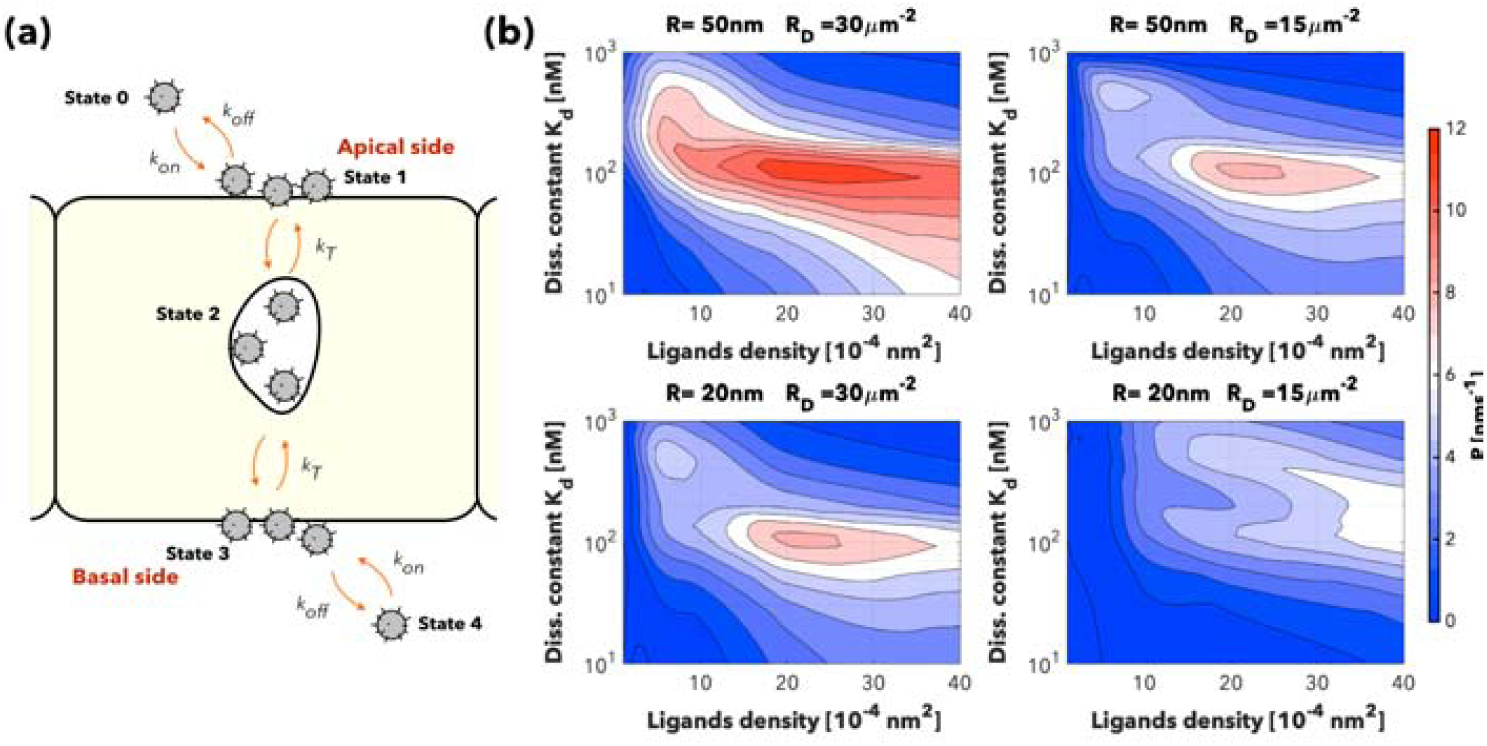
Modelling of the transcytosis of A_*L*_-POs. Representation of a theoretical model of transcytosis across endothelial cells showing five major steps: binding, endocytosis, trafficking, exocytosis and unbinding **(a)**. Apparent permeability (*P*) of POs plotted as function of both ligand number per particle (*L*) and the single ligand/receptor dissociation constant (*K*_*d*_) for nanoparticles with a radius *R* of 20 and 50 nm as well as receptor density *R*_*D*_ of 15 and 30 *μ*m−2.

**Figure S5:**
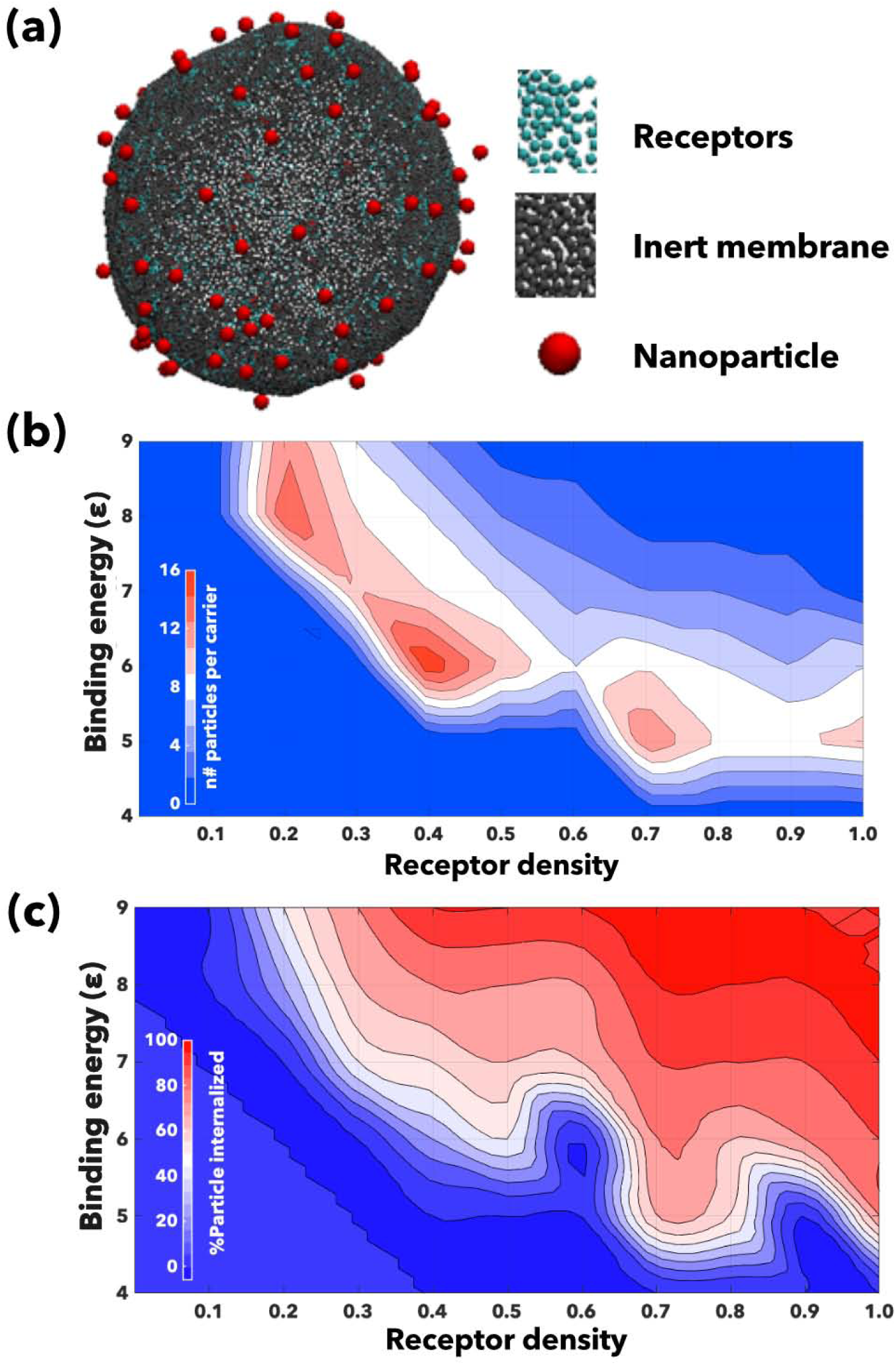
Molecular dynamics simulations of transcytosis of A_*L*_-POs. Simulations of the effect of avidity on membrane topological changes and nanoparticle aggregation dynamics. Coarse-grained membrane model using in the molecular dynamics simulations. Different beads were used to represent a membrane surface patch with no receptors (‘inert bead’ in black) and membrane surface patch with receptors (‘receptor bead’ in cyan). Nanoparticles are represented in red **(a)**. Number of particles per carrier **(b)** and percentage of particle internalised **(c)** plotted as function of binding energy and receptor density.

**Figure S6:**
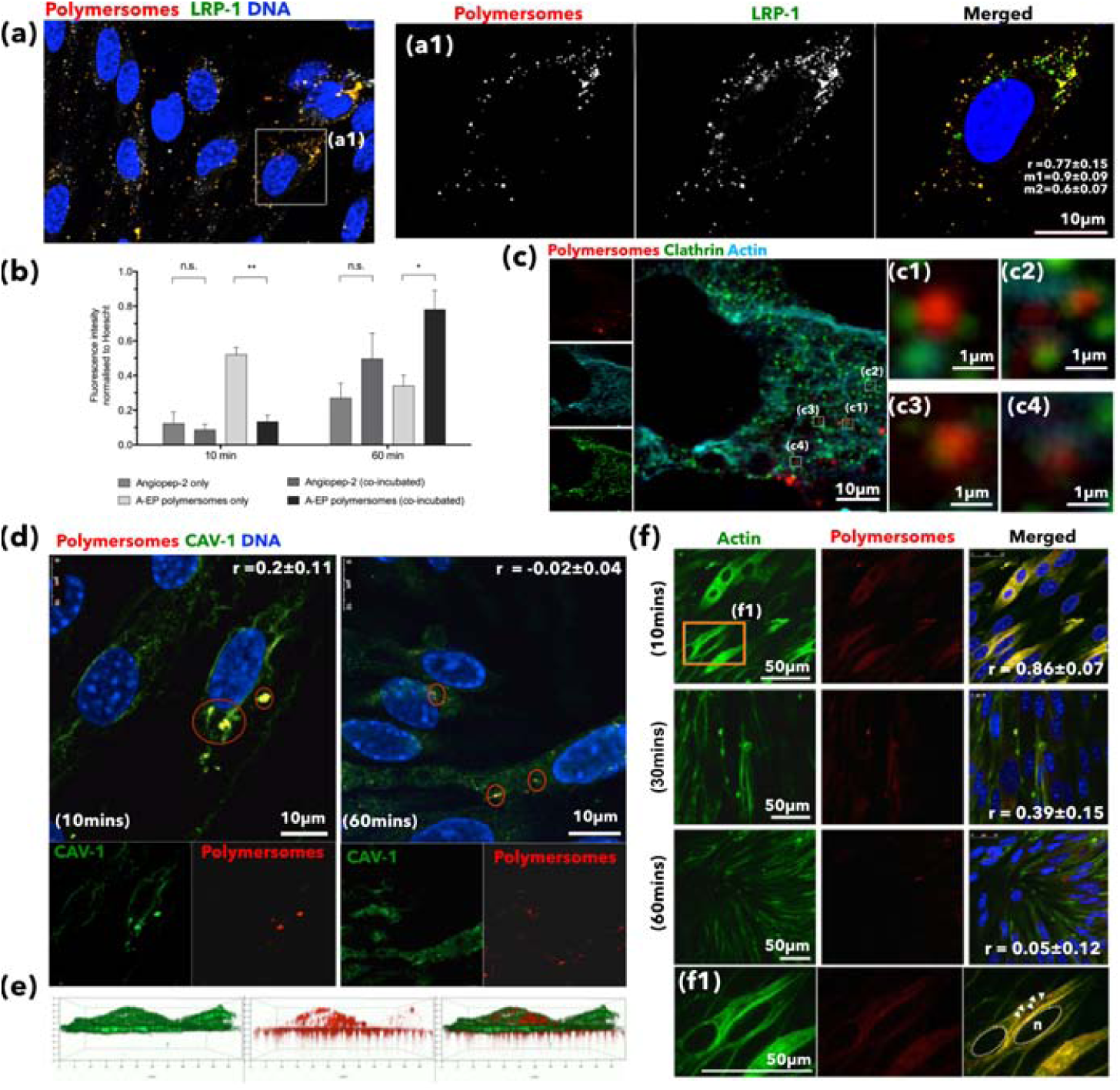
Transcytosis mechanism of A_22_-P across brain endothelial cells. Confocal micrograph of polarized bEnd3 showing colocalisation of Cy5-labelled A_22_-P (in grey) and LRP1 (in yellow) after 60 minutes of incubation **(a)**. Cell nuclei were counterstained with DAPI (in blue). Higher magnification of the co-localised Cy5-labelled A_22_-P and LRP1 on bEnd3 displaying the colocalisation values, *r* **(a1)**. Competition assay of angiopep-2 and A_22_-P. Comparison of intracellular fluorescence intensity of angiopep-2 or A_22_-P when incubated alone or co-incubated with polarised bEnd3. Data represented as mean ± SD (*n* = 3). * *P* <0.05 and ** *P* <0.01 **(b)**. Confocal image of clathrin (in green), Cy5-labelled A_22_-P (in red) and actin (in cyan) in bEnd3 treated with Cy5-labelled A_22_-P **(c)**. Higher manification of clathrin (in green) and Cy5-labelled A_22_-P (in red) staining on bEnd3 showing a partial colocalisation **(c1-c4)**. Confocal images of polarised bEnd3 incubated with Cy5-labelled A_22_-P (in red) for 10 and 60 minutes and stained for caveolin-1 (in green). Cell nuclei is stained with DAPI (blue). Colocalisation values *r* are displayed on the top corner of each image **(d)**. 3D rendering of bEnd3 incubated with Cy5-labelled A_22_-P (in red) and stained for caveolin-1 (green) **(e)**. Confocal micrographs of polarised bEnd3 stained for F-actin (Phalloidin in green) and incubated with Cy5-labelled A_22_-P (in red) for 10, 30 and 60 minutes. Nuclei stained with DAPI (blue). Colocalisation values *r* for F-actin and Cy5-labelled A_22_-P shown at the bottom of each merged image **(f)**. A higher magnification image showing colocalisation of Cy5-labelled A_22_-P and F-actin **(f1)**.

**Figure S7:**
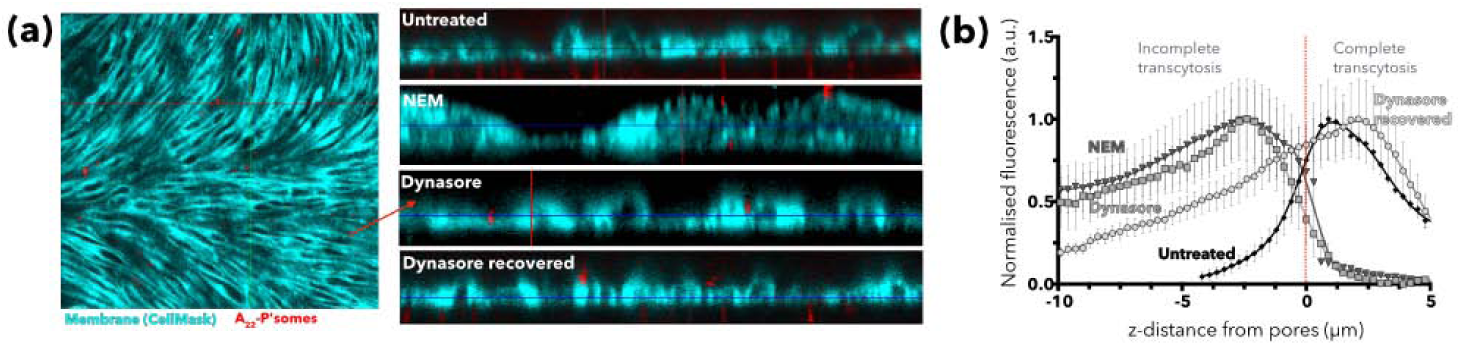
Dynasore and N-ethylmaleimide alter the transcytosis of A_*L*_-POs across brain endothelial cells. 3D rendering of polarised endothelial cells pre-treated with dynasore or N-ethylmaleimide (NEM) and further incubated for 60 minutes with Cy5-labelled A_22_-P (in red). Cell membrane is represented in cyan by staining with CellMask*™* **(a)**. Quantification of Cy5-labelled A_22_-P fluorescence intensity across confocal z-stack images of endothelial cells before and after pre-treatment with dynasore for 10 minutes. Dynasore recovery condition represents the cells that were washed with PBS after the incubation with dynasore. In the graph, zero represents the beginning of the pores, negative values above the transwell membrane and positive values within the transwell membreane. Data is represented as mean ± SD (*n* = 3) **(b)**.

**Figure S8:**
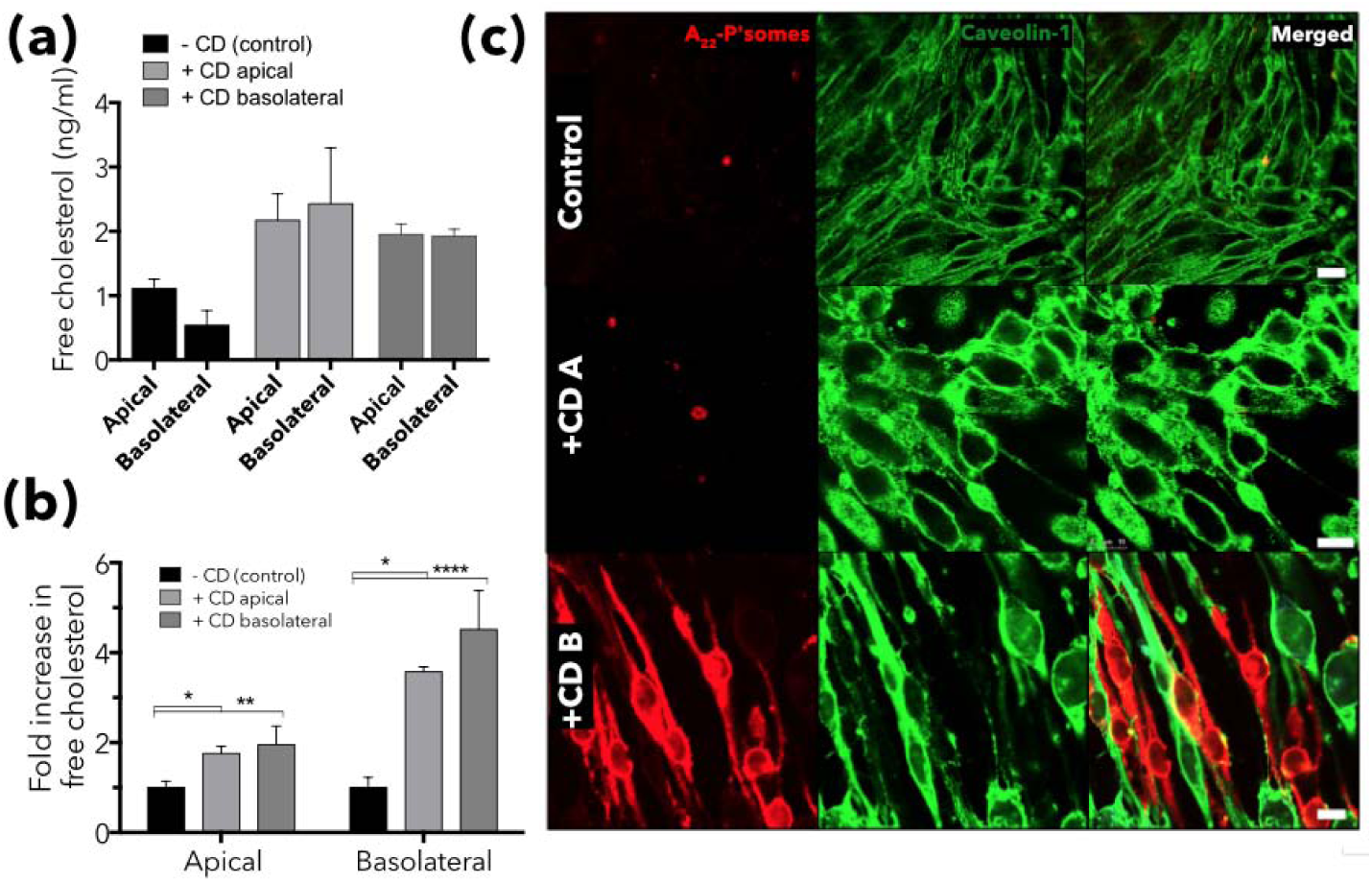
Cholesterol depletion impairs transcytosis of A_22_-POs across brain endothelial cells. Quantification of free cholesterol released into the media after depletion from the plasma membrane by incubation with methyl-*β*-cyclodextrin (CD) in the apical or basal side of the transwell. Data is represented as mean ± SD (*n* = 3) **(a)**. Fold-change in the level of cholesterol in the apical and basal side of the transwell membrane after incubation with CD. Data is represented as mean ± SD (*n* = 3). * *P* <0.05, ** *P* <0.01, **** *P* <0.0001 **(b)**. Confocal images of Cy5-labelled A_22_-P (red) in cholesterol-depleted bEnd3 cells either treated with CD in the apical (+CD A) or basal (+CD BL) side of transwell. Caveolin-1 is represented in green. Scale bar: 5*μ*m **(c)**.

**Figure S9:**
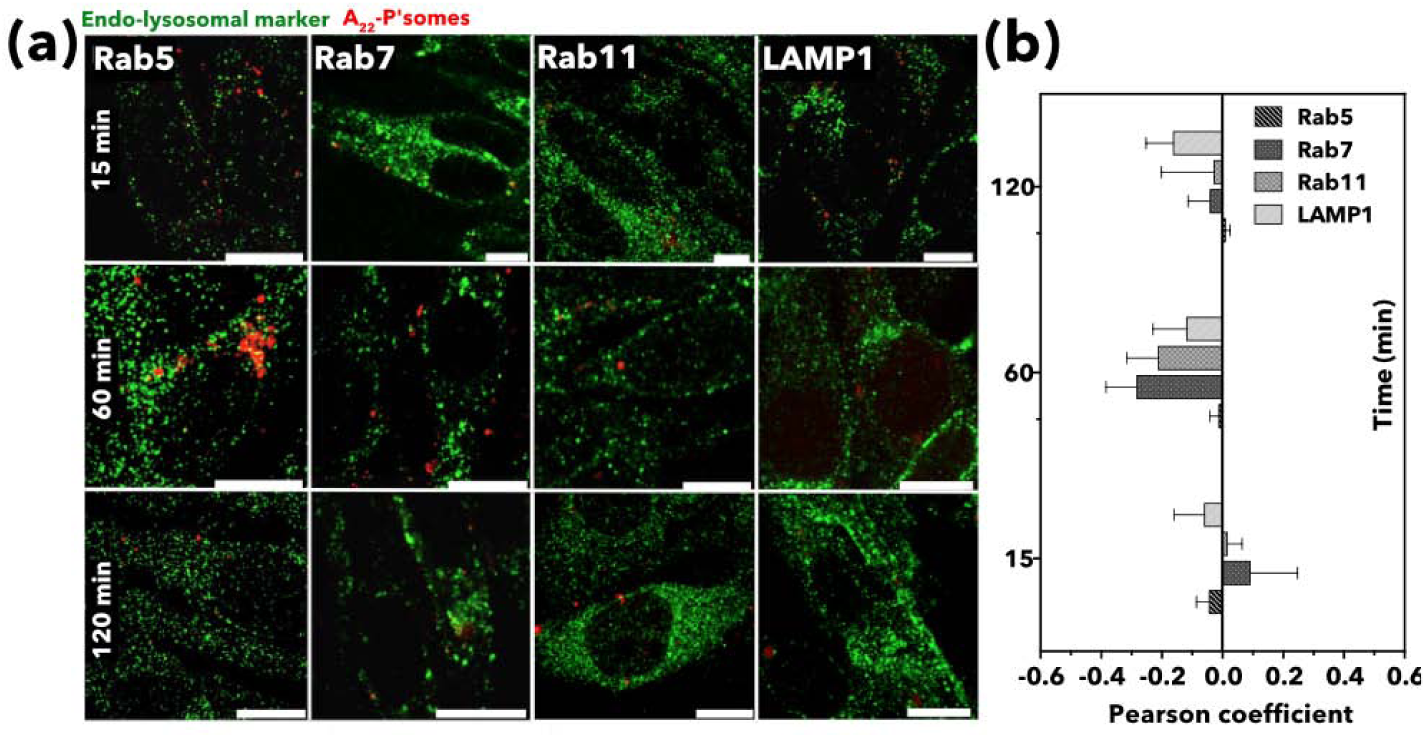
Endo- and lysosomal independent transcytosis of A_22_-POs across brain endothelial cells. Confocal images of Cy5-labelled A_22_-P (red) and markers of endosomal (Rab5, Rab7 and Rab11) and lysosomal (LAMP-1) (in green) endocytic pathways after incubation for 15, 60 and 120 minutes. Scale bar: 10 *μ*m **(a)**. Quantification of the colocalisation of Cy5-labelled A_22_-P with endosomal and lysosomal markers along the time **(b)**.

**Figure S10:**
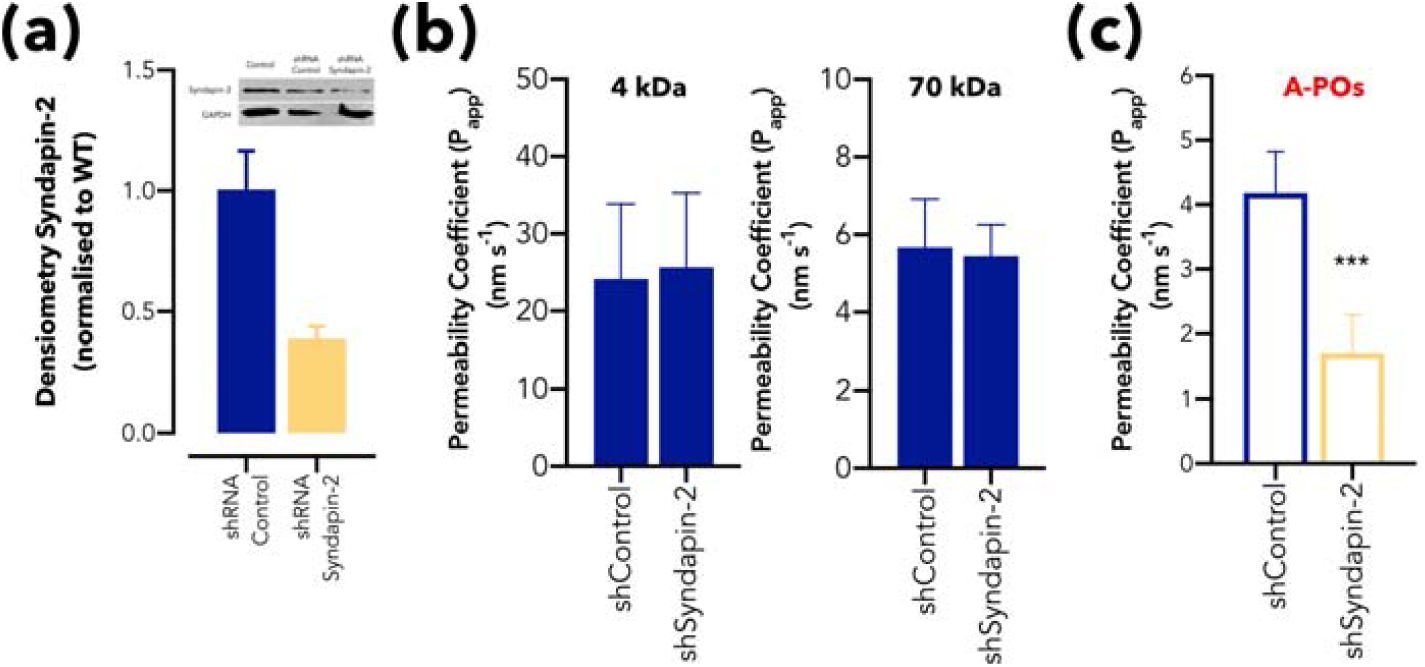
Syndapin-2 knockdown disrupts transcytosis of A_22_-POs. Expression of syndapin-2 on shRNA-transfected bEnd3 cells quantified by western blot. Levels of syndapin-2 on bEnd3 either treated with shRNA control or syndapin-2 were normalised to wild-type cells. Data is represented as mean ± SD (*n* = 6) **(a)**. Bar charts showing apparent permeability of 4 and 70 kDa dextrans across bEnd3 transfected with shRNA control or shRNA for syndapin-2 knockdown. Data represented as mean ± SD (*n* = 6) **(b)**. Apparent permeability of Cy7-labelled A_22_-P across control and syndapin-2 knockout bEnd3 cells. Data represented as mean ± SD (*n* = 6). *** *P* <0.001 **(c)**.

**Figure S11:**
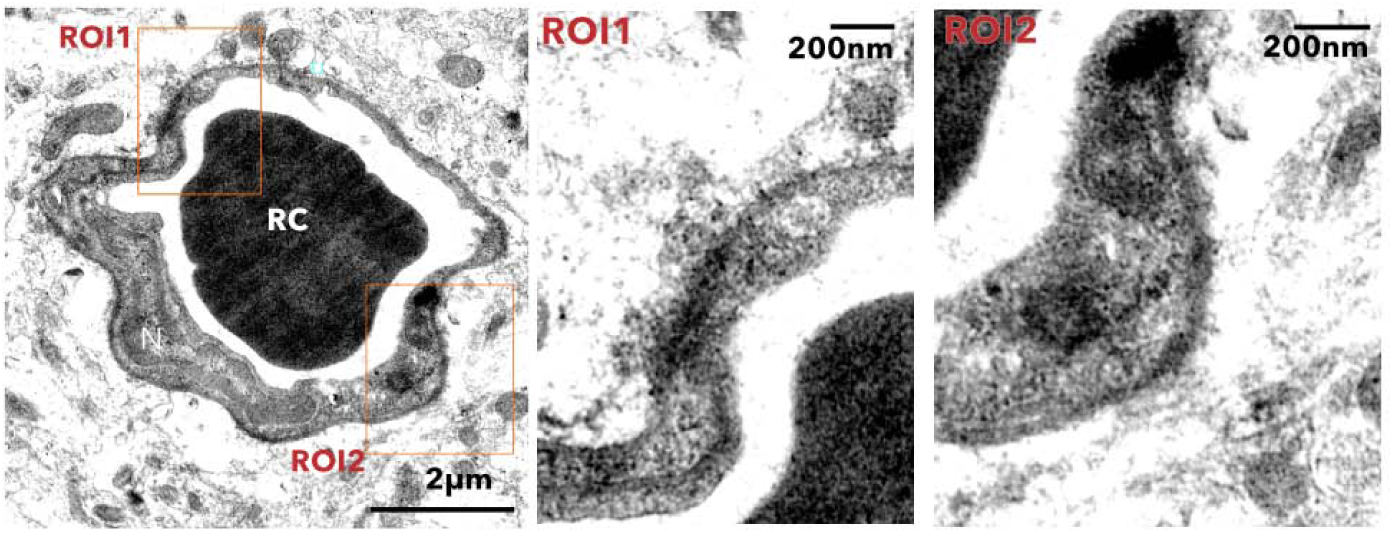
Penetration of PtA2-loaded A_22_-POs across *in vivo* BBB. Transmission electron micrograph of PtA2-loaded A_22_-P penetrating brain capillaries. Two high magnification images show in detail two regions of interest (ROI). RC: red blood cell and N: nucleus.

**Figure S12:**
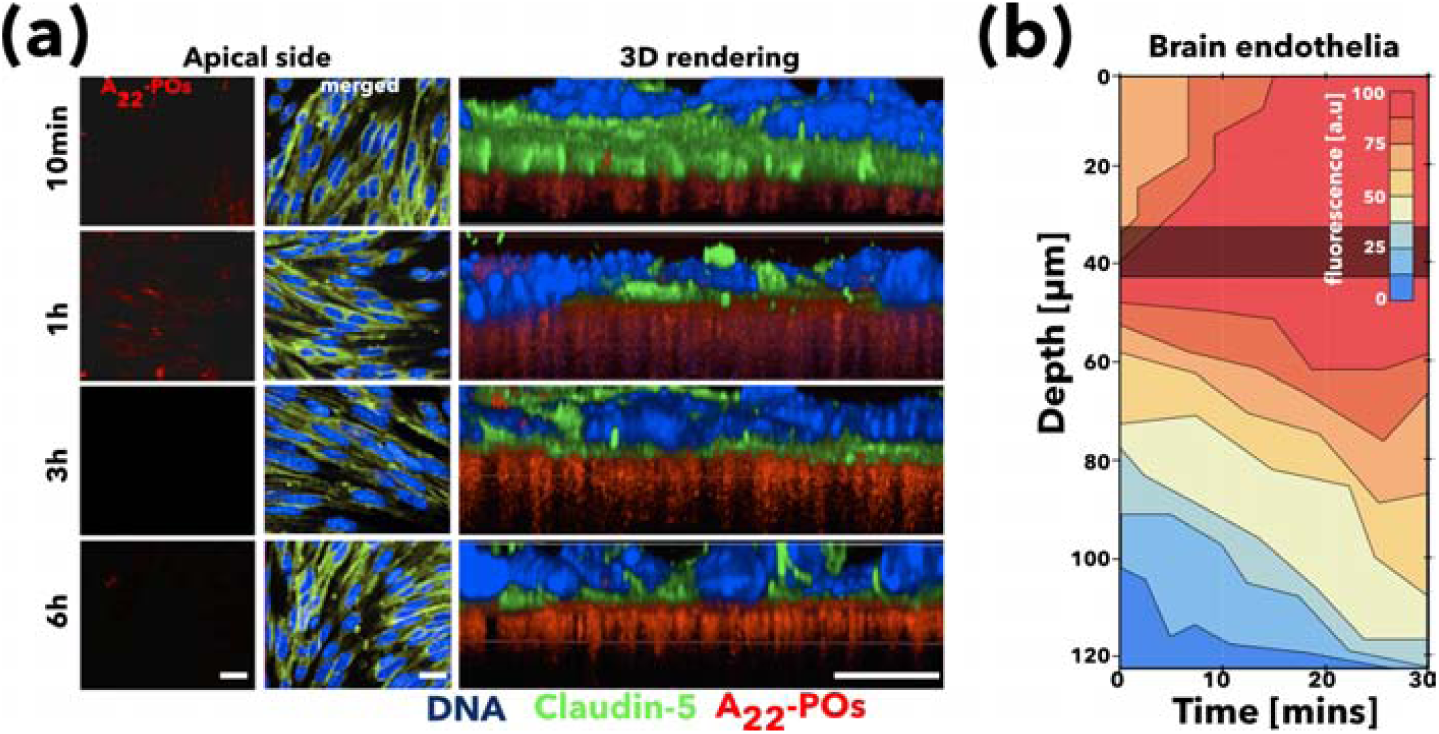
Live-cell imaging of A_22_-POs across brain endothelial cells. Confocal images and corresponding 3D rendering of polarised bEnd3 incubated with Cy5-labelled A_22_-P (red) for 10 minutes, 1, 3 and 6 hours. Tight junction, claudin-5, is represented in green and the nuclei counterstained with DAPI (in blue). Scale bar: 25 *μ*m **(a)**. Real-time 4D confocal imaging of transcytosis of Cy5-labelled A_22_-P. Heat map of Cy5-labelled A_22_-P fluorescence in the extended transwell area over time obtained through 4D imaging **(b)**.

